# Brain-wide microstrokes affect the stability of memory circuits in the hippocampus

**DOI:** 10.1101/2024.09.17.612757

**Authors:** Hendrik Heiser, Filippo Kiessler, Adrian Roggenbach, Victor Ibanez, Martin Wieckhorst, Fritjof Helmchen, Julijana Gjorgjieva, Anna-Sophia Wahl

## Abstract

Cognitive deficits affect over 70% of stroke survivors, yet the mechanisms by which multiple small ischemic events contribute to cognitive decline remain poorly understood. In this study, we employed chronic two-photon calcium imaging to longitudinally track the fate of individual neurons in the hippocampus of mice navigating a virtual reality environment, both before and after inducing brain-wide microstrokes. Our findings reveal that, under normal conditions, hippocampal neurons exhibit varying degrees of stability in their spatial memory coding. However, microstrokes disrupted this functional network architecture, leading to cognitive impairments. Notably, the preservation of stable coding place cells, along with the stability, precision, and persistence of the hippocampal network, was strongly predictive of cognitive outcomes. Mice with more synchronously active place cells near important locations demonstrated recovery from cognitive impairment. This study uncovers critical cellular responses and network alterations following brain injury, providing a foundation for novel therapeutic strategies preventing cognitive decline.

## Introduction

Although many stroke survivors develop forms of cognitive decline^1,2^, the pathomechanisms how cognitive decline emerges, even if cognitive brain areas are not directly affected, are not understood, nor are there any specific treatment options available. In particular, the accumulation of multiple, smaller ischemic events - with other obvious symptoms of stroke, such as motor impairment lacking - does not lead to acute cognitive impairment, but cognitive decline develops in months and years after the ischemic events^3^. The hippocampus, a relay station for cognition and memory processing, is particularly susceptible to ischemic events^4–6^, with neuronal death occurring already after a brief episode of ischemia^7,8^. Although this vulnerability of the hippocampus has been discussed in the context of its high plasticity, the anatomical connection to many brain areas and its vascularization^8,9^, it is not understood how individual neurons and functional networks in the hippocampus react to brain-wide injuries and how they rewire and recode to maintain their function. Several types of neurons coding for distinct memory functions have been identified: ÒKeefe and Dostrovsky^10^ discovered ‘place cells’ (PCs) in 1971 which respond specifically to the current location of the animal, but have been also discussed^11^ to contain compressed representation of contextual, sensory and episodic experiences during exploration of an environment. While the importance of ‘place cells’^10^ for memory formation and maintenance has been extensively demonstrated, it remains elusive, how place cells react individually or in ensembles to the induction of multiple microstrokes distributed throughout the brain. Revealing the functional impact of microstrokes on the single-cell level and on network tuning properties is of high translational value to better link neuropathological features after ischemic events to cognitive decline and to identify new targets for treatment strategies preventing cognitive deficits.

Here, we present a novel approach, where we studied the effect of brain-wide microstrokes on individual neurons and functional networks in CA1 of the hippocampus in mice performing a cognitive task. Using chronic two-photon calcium imaging in the hippocampus in mice navigating in a virtual reality corridor we could follow the fate of individual neurons in the healthy condition but also several weeks after the induction of disseminated cerebral microstrokes. We reveal that brain-wide microstrokes disrupt individual neuronal coding, functional hippocampal network architecture and induce cognitive decline. Our findings highlight the importance of understanding fundamental reorganization principles in the hippocampus on a cellular resolution level in relation to measurable cognitive deficit parameters: Analyzing the cellular and network responses after stroke, we could show that the cognitive outcome of animals is critically related to the individual stability of place cells or sub-networks of active surviving neurons with similar spatial tuning to maintain memory function and prevent cognitive decline.

## Results

### Microstrokes impair spatial memory and place cell stability

To identify the function of individual neurons during a spatial navigation task, we performed chronic two-photon calcium imaging of the same hippocampal network in the healthy condition (Figure 1A) and after stroke (Figure 1B). Mice expressing the calcium indicator GCaMP6f in CA1 were trained to navigate head-fixed in a virtual reality (VR) corridor (Figure 1A; Methods), while we simultaneously recorded calcium signals of neuronal populations in CA1 through a chronically implanted glass window 5 days before and up to 28 days after stroke surgery (Figure 1C, D).

**Figure 1:**
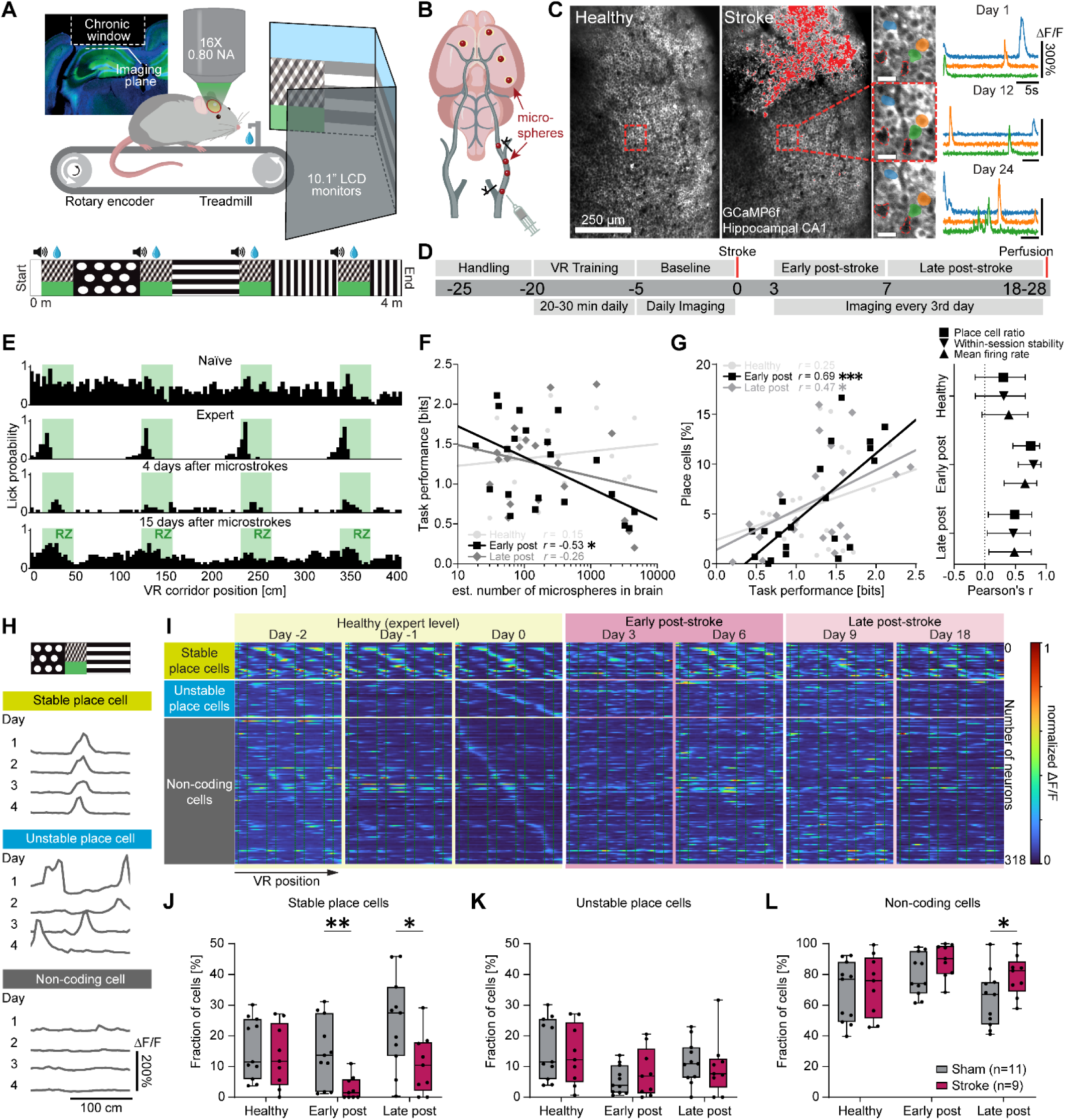
Microstrokes impair spatial memory and place cell stability. **A.** Experimental setup: Mice were trained to navigate in a linear virtual reality (VR) corridor (4 m length) during simultaneous unilateral two-photon calcium imaging in CA1 of the hippocampus. **B.** Schematic illustration of microstroke induction by injection of fluorescent microspheres (20 µm diameter) into the common carotid artery, inducing microstrokes. **C.** Representative two-photon images of the same field of view (FOV) in CA1 showing GCaMP6f fluorescence before (left) and after (middle) stroke induction. Damaged tissue overexposed the sensor and is shaded red. Individual neurons and their ΔF/F calcium fluorescence traces could be tracked over several weeks (right). Landmark blood vessels are marked in red. Scale bars = 25 µm. **D.** Timeline showing the sequence of events of the experiment in days relative to stroke induction. Chronic two-photon imaging in CA1 was performed while animals were navigating in the VR corridor during three experimental phases: before stroke (“healthy”, ≥-5 to 0 days before stroke), early (0-7 days) and late (>7 days – 28 days) after stroke. **E.** Histograms depicting the profile of an example mouse to lick for a water reward in the VR corridor. Green areas indicate reward zone locations. **F.** Number of microspheres in the brain plotted against the task performance (spatial information of lick profile, in bits) during the three phases. Inset shows Pearson’s correlation coefficients, data points are individual animals in each stroke phase. **G.** Left: Like F, but task performance plotted against the percentage of all imaged cells identified as place cells (place cell ratio). Right: Pearson’s correlation coefficients of task performance with metrics of neural activity such as the place cell ratio, the stability of neural spatial activity maps across trials (within-session stability) and the mean firing rate for the three phases. Error bars represent 95% confidence intervals. **H.** Spatial activity maps of three exemplary neurons being active at the same corridor location (stable place cell), different locations (unstable place cell), or at no specific location (non-coding cell) during four healthy sessions. **I.** Spatial activity maps of neurons imaged on multiple days throughout the experiment and sorted into the three functional classes from H (each row represents an individual neuron tracked before and after stroke). **J – L.** Percentages of all tracked cells that were classified as stable place cells (J), unstable place cells (K) and non-coding cells (L). In J – L, statistics were evaluated using two-way repeated-measures ANOVA with the Greenhouse-Geisser correction and Tukey-Kramer multiple comparisons test. Asterisks indicate significances: *p<0.05, **p<0.01, ***p<0.001.

We induced microstrokes by injecting fluorescent microspheres (20 µm diameter) unilaterally in the internal carotid artery, inducing disseminated microstrokes (Figure 1B, Figure S1). Microspheres were found directly visible under the hippocampal window (Figure S1A) in 4 out of 20 animals. We performed histological analysis to identify the microsphere distribution in the mouse brains and to quantify the lesion volume: Lesion volume and microsphere load were strongly correlated (Spearman’s ρ=0.84, p<0.001; linear regression model: slope = 0.001, R^2^=0.94; Figure S1D). Most microspheres and lesions were located in the neocortex (spheres: 39.3±2.2%, lesions: 31.3±4.3%), and in the hippocampus (spheres: 13.8±1.7%, lesions: 14.2±3.5%; Figure S1E), but also subcortically, e.g. in the thalamus and striatum (Figure S1E).

We trained mice to correctly identify four reward zones in the VR track where they received a water reward (Figure 1E). While naïve mice searched for water rewards in the entire corridor, expert mice learned to lick for water primarily in reward zones (> 60% licks within reward zones, Figure 1E). Microstrokes disrupted this licking pattern so that animals again randomly licked throughout the track, suggesting that mice lost the learned ability to locate themselves in the corridor. The more microspheres we found post-mortem in the brain, the worse was the performance of the animals in the VR corridor, particularly early (within 7 days) after stroke (r=-0.53, p=0.01; Pearson correlation, Figure 1F). Similarly, after induction of microstrokes animals with poor task performance displayed lower mean firing rates of CA1 neurons, a lower place cell ratio (percentage of all imaged cells that were classified as place cells, see methods), and a reduced stability of place cells to remain place cells across trials in individual sessions (“within-session stability”, Figure 1G). Animals with microstrokes displayed only transient, very minor neurological deficits on day 2 after stroke (Figure S2A, B), whereas we observed no impairment of task-relevant motor abilities (locomotion, Figure S2C, D; or licking, Figure S2E) compared to sham animals. A generalized linear model revealed no significant association between the number of microspheres in a distinct brain region and the cognitive performance in the VR corridor early after stroke (Figure S1F), suggesting that the number of accumulated microstrokes and thus the affected brain volume was more critical for the cognitive performance than the location of individual microstrokes.

Our experimental setup allowed us to track the activity of the same individual neurons before and after microstrokes for several weeks while animals performed the spatial navigation task. We could thus classify neurons according to the stability of their spatial tuning: We identified stable place cells, which were active when the mouse was at the same position in the VR during 5 consecutive days pre-stroke, whereas unstable place cells remapped their spatial field (Figure 1H, I). We could also identify non-coding cells, which did not meet the place cell criteria^12^ within imaging sessions (Figure 1H, I). When we compared the fractions of these three functional classes across experimental phases (healthy pre-stroke, early and late after stroke) and between stroke and sham animals (Figure 1J-L), we found that animals with microstrokes had significantly fewer stable place cells, both early and late after stroke compared to sham animals (fraction of stable place cells early after stroke: Sham: 14.7±3.3% versus Stroke: 2.9±1.3%, p=0.005; fraction of stable place cells late after stroke: Sham: 24.4±4.5% versus Stroke: 10.8±3.2%, p=0.025; Figure 1J). Stroke mice also had significantly more non-coding cells than sham animals (fraction of non-coding cells late after stroke: Sham: 64.1±5.5% versus Stroke: 79.3±4.4%, p=0.046; Figure 1L), while the fraction of unstable place cells remained unchanged for both groups (Figure 1K), indicating that microstrokes affected the functional properties of surviving neurons and the stability of spatial tuning.

### Microstrokes disrupt imprinting of distinct functional cell classes

As we had identified a loss of stable place cells after stroke, we next investigated whether individual surviving neurons had maintained or switched their functional class for spatial tuning over time from the healthy to the disease state (Figure 2A). As functional remapping of hippocampal networks is known to be naturally highly dynamic^13–15^, we first quantified the probabilities of place cells and non-coding cells to be a place cell on the next day in healthy networks, and compared these to chance level, simulated by a distribution where cell classes were randomly shuffled. We found that place cells were more likely to remain place cells than chance level, and than non-coding cells becoming place cells (Observed place cell to place cell (PC-PC) transitions: 28.5±3.4%; shuffled PC-PC transitions: 15.1±2.7%, versus observed PC-PC transitions: p<0.001; observed non-coding cell to place cell transitions (NC-PC): 13.5±2.3%, versus observed PC-PC transitions: p<0.001; Figure 2B). We term this phenomenon in CA1 neurons “functional imprinting”: Once the animal has learned the task, the corresponding neuronal network consolidates, meaning that neurons were attributed to distinct functions within the network making it less likely that these neurons randomly switched their dedicated function. We found that microstrokes disrupt this functional imprinting (comparing true and shuffled place cell transitions during post-stroke sessions between experimental groups, Figure 2C): While in sham animals the functional imprinting of a place cell to stay a place cell was maintained throughout the experiment (Figure 2C), the probability of place cells to stay place cells in stroke mice was at the same level as non-coding cells becoming place cells (Observed transitions in Stroke group: PC-PC transitions: 11.5±3.1% versus NC-PC transitions: 7.5±2.4%, p=0.400; Figure 2C left), and significantly lower than in sham mice in particular early after stroke (PC-PC transitions: Sham group: 27.1±4.3%, versus Stroke group: p=0.001; Figure 2C left), indicating that neurons are randomly assigned to different functions in the early phase after stroke. Additionally, the probability of place cells to stay place cells the next day was not significantly different from a shuffled distribution anymore (shuffled PC-PC transitions in Stroke group: 8.8±2.7%, versus observed PC-PC transitions in Stroke group: p=0.691; Figure 2C, left). Notably, this effect was only transient in the early phase after stroke and recovered within 3-4 weeks after stroke surgery (Figure 2C right).

**Figure 2:**
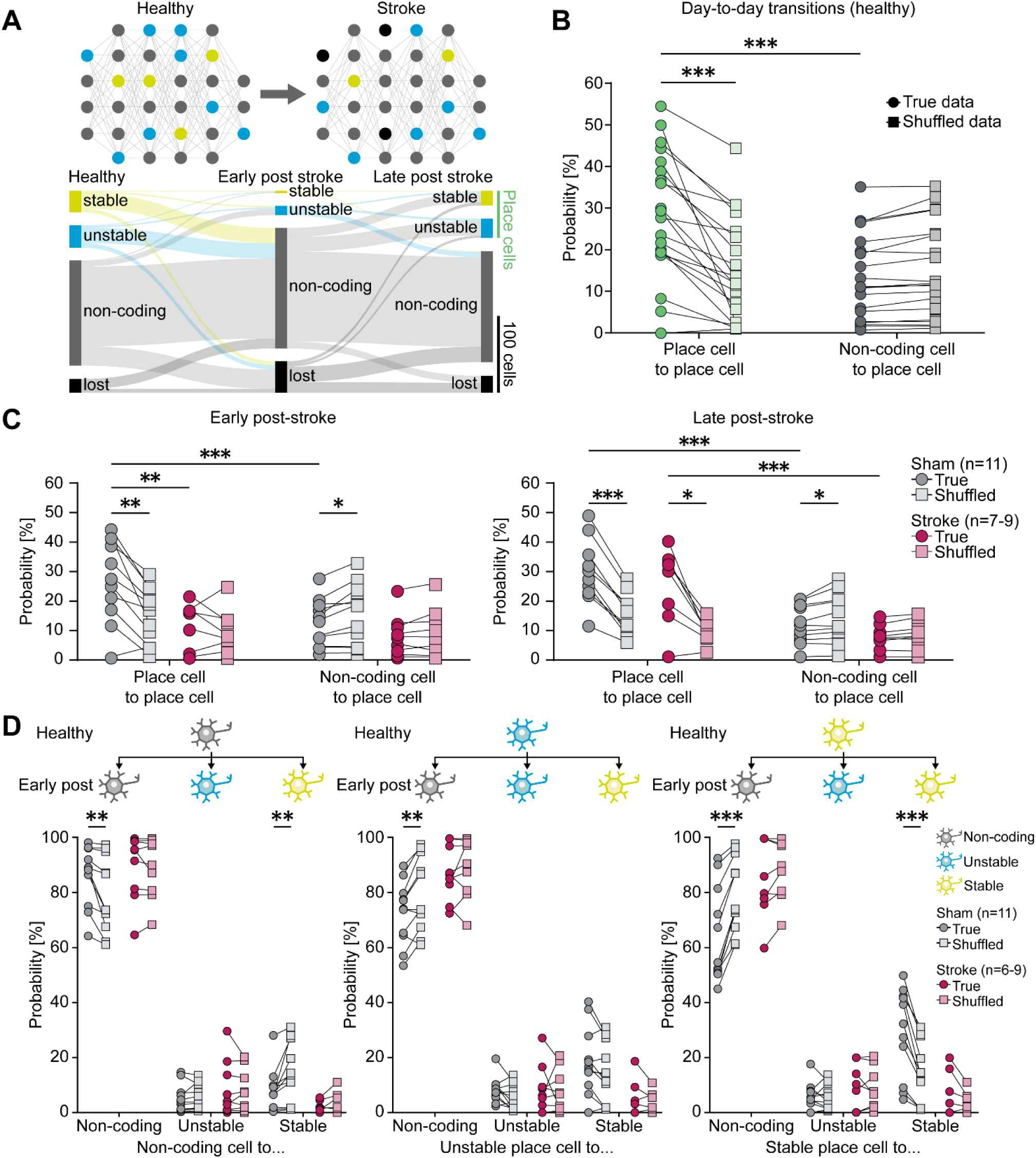
Microstrokes disrupt imprinting of distinct functional cell classes. **A.** Top: Scheme reflecting how neurons in hippocampal networks can transition between functional classes, which can be affected by microstrokes. Bottom: Sankey diagram showing the number of neurons in all stroke mice maintaining or changing functional cell classes between stroke phases: Neurons can transition to and from being non-coding for spatial information (NCs, grey), place cells (PCs, green), either stably (yellow) and unstably (blue) coding for space, as well as lost cells (black) that could not be tracked in a given phase. Height of bars represent number of neurons in each class per phase. Connection width is the number of cells that switch functional class. **B.** The probability of place cells (green) or non-coding cells (grey) to become place cells (PCs) on the next days in healthy networks. Data in dark circles show observed transitions, data in light squares present transitions at chance level in distributions where functional classes were randomly shuffled. Subject-matched mixed-effects model with Bonferroni multiple comparisons test. **C.** Transition probabilities between experimental groups and shuffled distributions during post-stroke phases. **D.** Observed probabilities of non-coding cells (grey), unstable (blue) and stable (yellow) place cells to maintain or switch their functional class from the healthy to the early post-stroke phase. Two-way repeated-measures ANOVA with Bonferroni multiple comparisons test. Asterisks indicate significances: *p<0.05, **p<0.01, ***p<0.001.

We next examined if functional imprinting also occurs when comparing the healthy and post-stroke condition of the three identified functional cell classes (Figure 1H, I): non-coding cells, unstable and stable place cells. Indeed, in sham animals stable place cells (sPC, yellow) and non-coding cells (NC, grey) preferably maintained their function comparing observed transition probabilities to chance level (Sham group early post-stroke: observed NC-NC transitions: 86.5±3.3% versus shuffled NC-NC transitions: 79.7±4.1%, p=0.001; observed sPC-sPC transitions: 29.4±4.9% versus shuffled sPC-sPC transitions: 14.7±3.3%, p<0.001, Figure 2D). The transition to another functional class was less likely (Sham group early post-stroke: observed NC-sPC transitions: 8.3±2.3% versus shuffled NC-sPC transitions: 14.7±3.3%, p=0.003; observed sPC-NC transitions: 64.7±5.2% versus shuffled sPC-NC transitions: 79.7±4.4%, p<0.001). This was in strong contrast to stroke animals, where transition probabilities between all three classes were at chance level early after stroke (Stroke group early post-stroke: e.g. observed sPC-sPC transitions: 5.9±2.8% versus shuffled sPC-sPC transitions: 3.3±1.4%, p=0.703; observed sPC-NC transitions: 85.1±5.1% versus shuffled sPC-sPC transitions: 87.8±3.9%, p=0.637, Figure 2D). However, microstrokes only transiently disrupted the stability of spatial coding, as the functional imprinting for place cells recovered in the later phase after stroke (Stroke group late post-stroke: observed sPC-sPC transitions: 31.0±10.3% versus shuffled sPC-sPC transitions: 15.1±4.4%, p=0.051; Figure S3) with a lower probability of stable place cells becoming non-coding cells (Stroke group late post-stroke: observed sPC-NC transitions: 54.0±12.4% versus shuffled sPC-NC transitions: 71.0±4.7%, p=0.030, Figure S3). Interestingly, in the late phase of the experiment, sham animals showed a tendency of unstable place cells to turn into stable place cells indicative of a functional consolidation of the network for the spatial navigation task on long-term (Sham group late post-stroke: observed uPC-uPC transitions: 21.9±4.3% versus shuffled uPC-uPC transitions: 11.5±2.1%, p=0.023; observed uPC-NC transitions: 50.6±7.4% versus shuffled uPC-NC transitions: 64.0±5.6%, p=0.001, Figure S3).

### Functional stability of spatial coding influences cognitive outcome

Having identified the loss of stability of spatial coding in surviving neurons as a major effect after stroke, we next investigated how important the stability of spatial coding is for the cognitive outcome of the animals in the spatial navigation task. When comparing the post-stroke task performance to correctly identify reward zones in the VR corridor to the pre-stroke level, all stroke mice displayed a significant cognitive deficit 3 days after stroke (<75% below their healthy baseline, Figure 3A). However, while some animals recovered from the cognitive deficits within 10 days after stroke (“Recovery” group, Figure 3A), this was not the case for other animals that showed a chronic cognitive deficit (“No-Recovery” group, Figure 3A). Animals with this chronic cognitive deficit lost a significant fraction of place cells with stable place fields on long-term compared to sham animals (Stable place cells in late post-stroke: No-Recovery group: 5.5±2.8% versus Sham group: 24.4±4.5%, p=0.009, Figure 3B). Animals with a recovery of the cognitive deficit however, showed only transiently a reduced number of stable place cells early after stroke (“Recovery” group: 4.5±2.1% versus Sham group: 14.7±3.3%, p=0.050), with the number of stable place cells increasing again, late post-stroke with no significant difference from sham animals (Recovery group: 15.1±4.8%, versus Sham group: p=0.084). These results suggest that the maintenance of individual place cells to keep their preference for a place field is important for the cognitive outcome.

**Figure 3:**
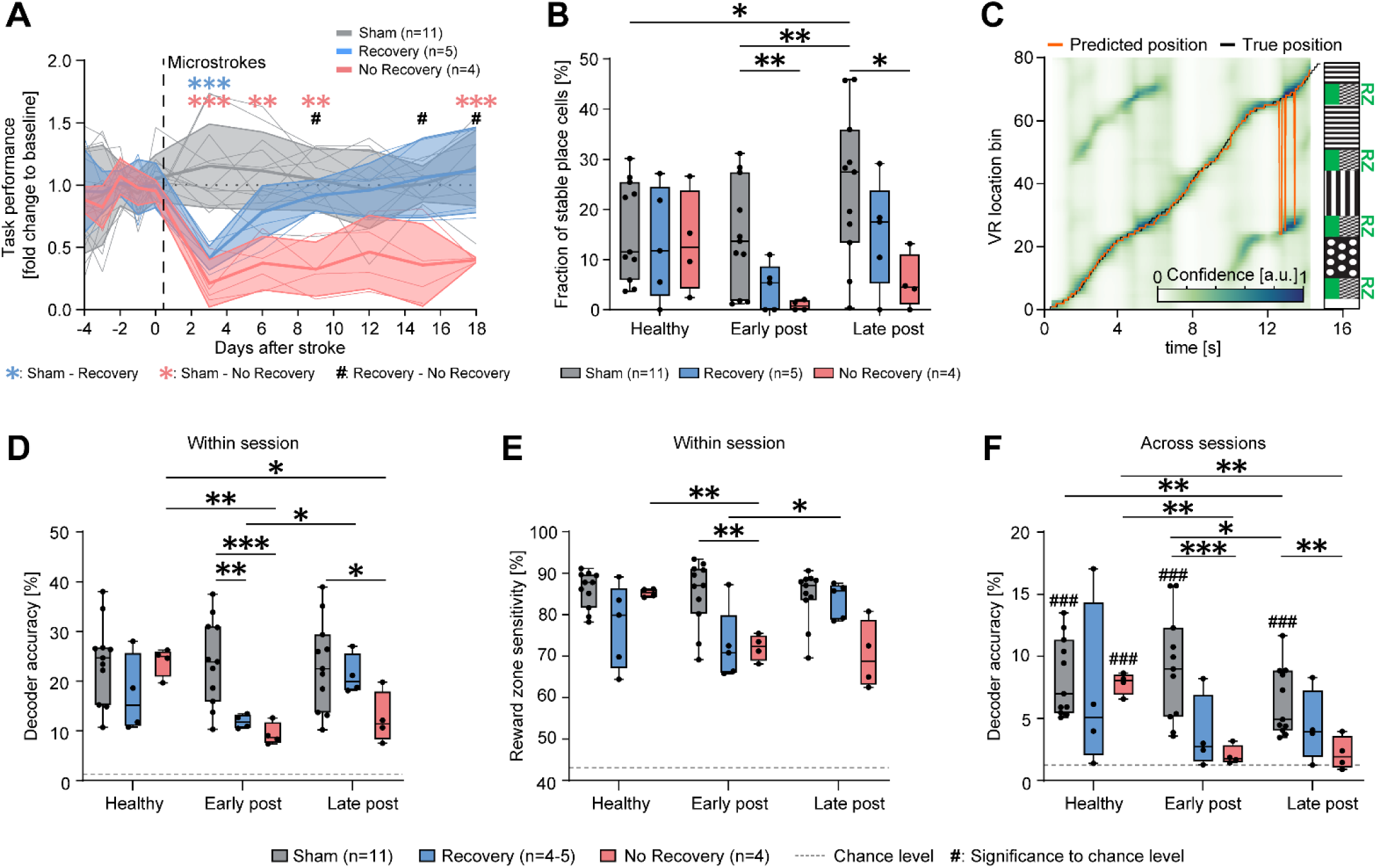
Functional stability of spatial coding influences cognitive outcome. **A.** VR performance after stroke relative to the healthy baseline revealing two different cognitive outcomes: A subset of mice showed a chronic cognitive deficit (“No-Recovery group” in red), while another group recovered from the initial cognitive decline within one week (“Recovery group” in blue). Sham-operated animals (grey) maintained their VR performance. A mixed-effects model with the Greenhouse-Geisser correction and Tukey-Kramer multiple comparisons test revealed significantly reduced task performances depending on the stroke outcome group (F_Group×Time_(22, 180)=5.380, p<0.001). Asterisks denote significance levels between sham and “Recovery” (blue) or “No-Recovery” (red) groups. Hash symbols indicate significance levels between “Recovery” and “No-Recovery” groups. **B.** Percentages of stable place cells out of all imaged neurons in the three outcome groups across experimental phases. **C.** Predicted corridor positions generated by a Bayesian decoder (orange) from neural ΔF/F traces and true position (grey) of a single exemplary trial. The background heatmap visualizes frame-wise decoder confidence (arbitrary units) for each VR position bin. The bin with the highest confidence is the prediction of the decoder for each frame. Prediction errors (e.g. around second 13) can often be attributed to the confusion of different reward zones with similar wall patterns. **D.** Accuracy (percentage of frames with correctly predicted position bin) of the decoder indicating performance on data from the same session on which the decoder was trained (within-session decoder). Dashed line indicates chance level (1.25%). Data from one mouse was excluded with a decoder accuracy not significantly different from chance level in the healthy condition. **E.** The sensitivity of the within-session decoder to detect reward zones (percentage of frames within reward zones correctly predicted as being in a reward zone). Dashed line indicates chance level (43.75%). **F.** Accuracy of the cross-session decoder (decoder trained on the last healthy session and applied to all other sessions). Hash symbols indicate significance (one-sample t-tests with Bonferroni correction) to chance level (dashed line, 1.25%). Group differences in B, D-F were evaluated with two-way repeated measures ANOVA with the Greenhouse-Geisser correction and Tukey-Kramer multiple comparisons tests. Asterisks and hash symbols indicate significances: */# p<0.05, **/## p<0.01, ***/### p<0.001.

We then examined if population activity coding can predict the position of the animal in the VR corridor before and after stroke^17–19^. To investigate whether microstrokes affect the quality and stability of this encoding, we applied a Bayesian decoder to predict the most likely corridor position at each frame from neural activity of the same session^16,20^ (Figure C). The decoder was able to predict the correct position (Figure D) and detect reward zones (Figure E) significantly above chance levels in all groups in the healthy condition. However, the accuracy of the decoder and its sensitivity to predict reward zones were significantly lower for both stroke groups compared to sham early post-stroke (Decoder accuracy, Figure 3D: Sham: 23.8±2.6%, Recovery: 11.8±0.7%, versus Sham: p=0.003; No-Recovery: 9.3±1.1%, versus Sham: p<0.001. Reward zone sensitivity, Figure 3E: Sham: 85.2±2.4%, Recovery: 72.5±3.9%, versus Sham: p=0.063; No-Recovery: 72.1±1.6%, versus Sham: p=0.002). The decoder showed a significantly improved accuracy and sensitivity to predict the animal’s position for the Recovery group in the late phase after stroke (Accuracy: late post-stroke: 21.2±2.0%, versus early post-stroke: p=0.038; Sensitivity: late post-stroke: 83.5±1.9%, versus early post-stroke: p=0.045), consistent with the recovery of the cognitive deficits in these mice.

To further understand the long-term stability of neural network encoding for spatial information we adapted the Bayesian decoder by training it on the last healthy session before stroke and applying it to all other sessions. This long-term decoder could predict the corridor positions of sham mice significantly above chance level throughout the entire experiment (Figure F). However, the ability of the long-term decoder to predict the position of animals based on their population activity was abolished for animals with micro-strokes and particularly for those with a chronic deficit (Figure 3F, Decoder accuracy in early post-stroke: Sham: 9.1±1.3% versus No-Recovery: 2.0±0.4%, p<0.001; late post-stroke: Sham: 6.4±0.8% versus No-Recovery: 2.2±0.7%, p=0.007).

### Stability, precision and persistence of functional network structure are markers for cognitive outcome

We next examined the stability of spatial memory on a network level. We performed population vector correlation (PVC)^16^, which quantifies the stability of the spatial activity of a network across days and can be visualized in cross-correlation matrices (Figure 4A top). Whereas sham animals showed a stable pattern of population activity relative to the position of the mice in the VR corridor throughout all stages of the experiment, animals in the No-Recovery group revealed a disturbed pattern compared to the healthy condition throughout both post-stroke phases (Figure 4A). In contrast, the pattern of population activity recovered 2-4 weeks after stroke for animals in the Recovery group, consistent with their cognitive recovery in the spatial navigation task (Figure 4A). Averaging the cross-correlation matrices across corridor location offsets yields summary curves that represent the similarity of neural activity at any two corridor locations with increasing distance (Figure bottom), allowing not only to quantify the stability of spatial information coding in neural populations across time, but also the spatial specificity and precision of that coding. In sham animals the matrices and curves showed clear periodic sequences corresponding to the layout of the VR corridor with repetitive reward zones and inter-reward zone areas with diverse wall patterns (Figure 4A). In contrast, periodicity was lost in No-Recovery mice with only flat population vector curves after microstrokes (Figure 4A), while periodicity re-emerged in animals of the Recovery group.

**Figure 4:**
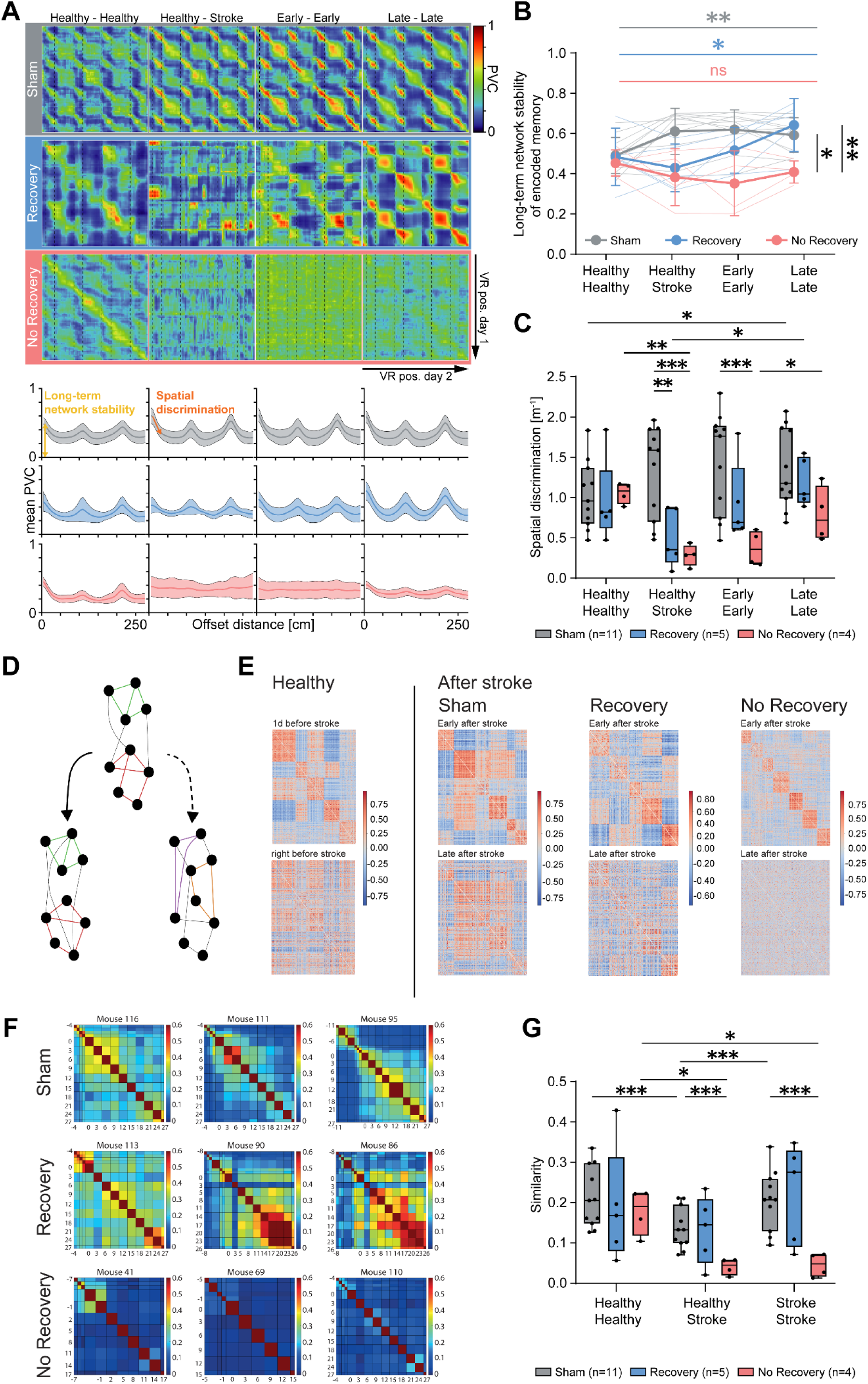
Stability, precision and persistance of functional network structure are markers for cognitive outcome. **A.** Top: Population vector correlation (PVC) matrices of outcome groups across session pairs within each experimental phase. “Healthy – Stroke’’ depicts matrices of the last healthy session in relation to the first post stroke session. Dashed lines show reward zone borders. Below: Average PVC curves summarize matrices across corridor location offsets. Correlation peaks reflect the periodic structure of the VR corridor. **B.** Y-intercept of PVC curves, which indicates cross-session stability of functional coding, in the three outcome groups across experimental phases. Thin lines represent individual mice, thick lines with error bars show mean and standard deviation. Horizontal significance bars mark time effects (group-wise one-way repeated-measures ANOVA), vertical significance bars mark differences between groups in the later phase after stroke (>7 days post injection). **C.** The absolute initial slope of PVC curves, which represents spatial precision in neural location coding, with higher values (steeper slopes) indicating higher precision. **D.** Change of functional network structure (correlations of the spatially binned activity) over time varies with effect of stroke. Schematic to represent functional network structure before (top) and after (bottom) surgery, with two possible outcomes: functional structure before stroke largely persists (left, solid arrow) or significantly changes (right, dashed arrow). **E.** Example matrices of functional correlations of spatially binned activity on subsequent experimental sessions. In healthy mice, functional structure on a given day (bottom) largely resembles the functional structure on the previous day. After stroke, sham mice and recovery mice exhibit similar functional structure of their spatial activity maps across consecutive days. Instead, No-Recovery mice show very different functional structure of their spatial activity maps even on subsequent days. **F.** Cosine similarity of functional correlations of spatially binned activity on different days (computed for the off-diagonal elements of the spatial correlation matrices shown in E). **G.** Mean similarity for different stroke groups. Each data point shows the mean of all similarities in F, when the sessions being compared are both before stroke (Healthy-Healthy), before and after stroke (Healthy-Stroke) and both after stroke (Stroke-Stroke). Group differences in B, C and G were evaluated with two-way repeated measures ANOVA with the Greenhouse-Geisser correction and Tukey-Kramer multiple comparisons tests. Asterisks indicate significances: *p<0.05, **p<0.01, ***p<0.001.

The y-intercept of the population vector curves represents the correlation of the population activity at the same corridor location on two different days. It can be interpreted as a measure of the cross-session stability of the encoded memory (long-term network stability, Figure 4A). When comparing the y-intercept between experimental groups and across time (Figure), we find that mice in the Sham and Recovery groups displayed a significant increase in cross-session stability of the neuronal network over time, suggesting a consolidation of network activity for spatial memory (y-intercept change over time: Sham group: F(1.4,14.1)=12.50, p=0.002; Recovery group: F(2.1,8.5)=6.94, p=0.016; Figure 4B). In contrast, animals with a chronic deficit did not show improved cross-session stability over time (No-Recovery group: F(2.1,6.4)=1.1, p=0.405), which instead remained on a significantly lower level late after stroke (y-intercept in late post-stroke: No-Recovery: 0.41±0.03; Sham: 0.59±0.03, versus No-Recovery: p=0.003; Recovery: 0.64±0.06, versus No-Recovery: p=0.030; Figure 4B)

Next, we quantified the ability of the neuronal networks recorded in CA1 to distinguish nearby corridor location as a measurement of the precision of spatial coding. The initial slopes of the mean PVC curves (Figure 4A) show the rate at which population activity becomes different when comparing different locations, with a steeper slope indicating a higher spatial discrimination and memory precision. We find that spatial discrimination is strongly disrupted in both stroke groups immediately after stroke compared to sham (Precision in the healthy versus the post-stroke network activity: Sham: 1.33±0.17; Recovery: 0.49±0.16, versus Sham: p=0.009; No-Recovery: 0.28±0.07, versus Sham: p<0.001; Figure). However, the deficits were not permanent. Although No-Recovery mice showed some improvement of spatial precision late post-stroke (early post-stroke: Sham: 1.44±0.19 versus No-Recovery: 0.37±0.11, p<0.001; late post-stroke: Sham: 1.35±0.15 versus No-Recovery: 0.79±0.17, p=0.087), Recovery mice displayed a faster and more complete re-establishment of spatial discrimination already within the first week post-stroke, similar to the sham group (early: Recovery: 0.93±0.23, versus Sham: p=0.242; late: Recovery 1.18±0.13, versus Sham: p=0.678; Figure 4C).

We also assessed the effect of microstrokes on the persistence of functional network structure in the hippocampus over time (Figure 4D). First, we computed the functional correlations of the spatially binned activity across pairs of neurons on each day (Figure 4E). We found that in healthy networks, the resulting functional networks looked similar, especially across consecutive days (“Healthy”, “Sham”, Figure 4E). While the same appeared to hold for Recovery mice when comparing networks after stroke (“Recovery”, Figure 4E), in No-Recovery mice no resemblance of functional correlation structure could be seen after stroke. To quantify the persistence of functional networks across time in the different experimental groups, we computed the cosine similarity of the functional correlation structure of tracked neurons on all pairs of days (Figure 4F). Indeed, while there were no differences across the three groups of mice before stroke induction, No-Recovery mice showed significantly lower functional network similarity after stroke compared to Sham (Stroke-Stroke: No-Recovery: 0.05±0.02, Sham: 0.21±0.02, p<0.001; Figure 4G). We also observed a lower similarity when comparing the functional networks of sham mice separated by many days (healthy-stroke) as opposed to consecutive days, consistent with a slight drift (Sham: Healthy-Stroke: 0.13±0.02; Healthy-Healthy: 0.21±0.02, vs. Healthy-Stroke: p<0.001; Stroke-Stroke: 0.21±0.02, vs. Healthy-Stroke: p<0.001; Figure 4G). The similarity between functional networks of No-Recovery mice separated by many days, however, was significantly lower than this “normal” ongoing drift (Healthy-Stroke: No-Recovery: 0.04±0.01, vs. Sham: p<0.001; Figure 4G).

Hence, in healthy mice, groups of neurons show strong functional correlations of their spatially binned activity that persist across time, demonstrating stability of functional subnetworks. Following stroke, stable functional subnetworks return only in sham and recovery mice (albeit different than before stroke), while in No-Recovery mice functional subnetworks continue changing across time, consistent with the circuit’s inability to recover normal behavior. Therefore, these data suggest that the stability of functional networks over time could be essential in maintaining cognitive capability after microstrokes.

### Synchronous activity close to salient locations in animals with recovery of memory deficits

To investigate a functional relationship between individual surviving neurons after stroke, we quantified the synchronicity of pairs of neurons by correlating their ΔF/F traces (Figure 5A). We found no significant differences on average for all pairs of neurons comparing the three outcome groups and the healthy and post-stroke phase (Figure 5B). However, when examining neuronal pairs with highly correlated activity (95^th^ percentile), we found that animals with chronic cognitive deficits (No-Recovery group) had significantly less synchronously active place cells early after stroke than sham animals (Sham: 18.4±2.1% vs. No-Recovery: 8.3±2.3%, p=0.048; Figure 5C), while after stroke the number of synchronous active place cells increased for the Recovery group (Early post-stroke: 15.9±3.0% vs. Late post-stroke: 24.5±3.6%, p=0.025; Figure 5C). Correspondingly, the number of synchronously active non-coding cells expanded in the No-Recovery group compared to Sham early and late after stroke (Early post-stroke: Sham: 4.4±0.1% vs. No-Recovery: 4.9±0.1%, p=0.015; Late post-stroke: Sham: 4.3±0.2% vs. No-Recovery: 4.8±0.1%, p=0.033; Figure 5D). When investigating the similarity of spatially binned activity in the 95^th^ percentile of highly correlated neuronal pairs, we found less neuronal pairs with similar spatial activity in the No-Recovery group compared to the healthy condition and the Sham group (No-Recovery: healthy 0.47±0.02 vs. early post-stroke: 0.38±0.02, p=0.027; Sham (early post-stroke): 0.51±0.01 vs. No-Recovery (early post-stroke): p=0.001; Figure 5E). In contrast, neuronal pairs with similar spatial activity re-emerged after an initial drop early after stroke in animals with a recovery of the memory deficit (Recovery group: early post-stroke: 0.47±0.03 vs. late post-stroke: 0.56±0.03, p=0.021; Figure 5E).

**Figure 5:**
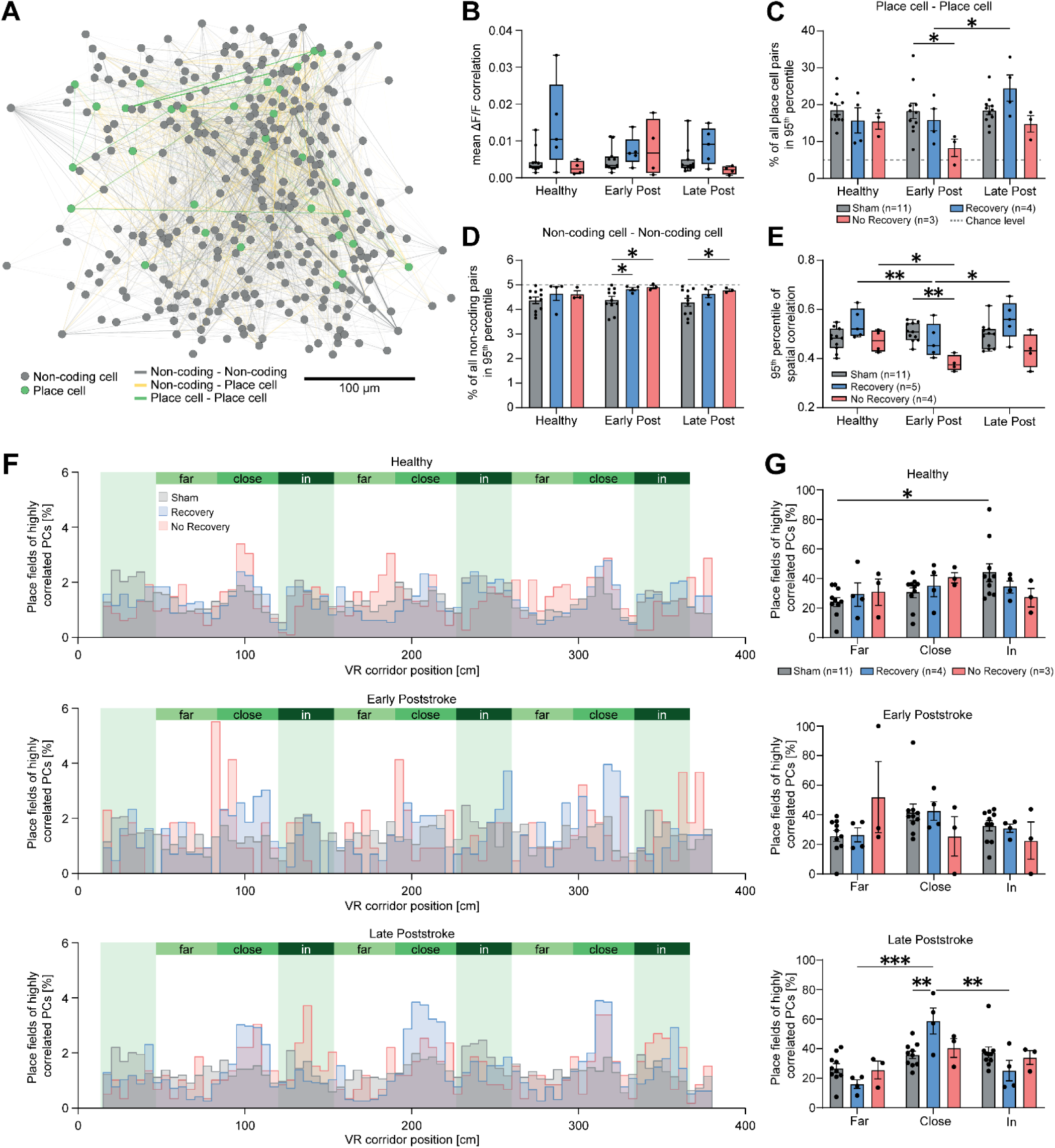
Synchronous activity close to salient locations in animals with recovery of memory deficits. **A.** Network schematic showing spatial distribution and functional connectivity (FC, Pearson’s correlation of ΔF/F traces) between all imaged place cells (PCs, green) and non-coding cells (NCs, gray) in an exemplary field of view. Highly correlated neurons (95th percentile) are connected by lines. Line width and opacity represent connectivity strength, and colors identify functional coding of cell pairs (gray: NC – NC pairs; yellow: NC – PC pairs, green: PC – PC pairs). **B.** Average FC between all imaged neurons across experimental groups. **C.** Percentage of place cell pairs in the 95^th^ pool of cells with highly correlated activity shows a loss of place cells pairs in No-Recovery group while at the same time more highly correlated non-coding cells were detected in the No-Recovery group (**D).** Error bars depict standard error. Dashed lines represent chance level (5%)**. E.** 95th percentile of correlation coefficients of spatial activity maps between all imaged neurons. **F.** Histograms of place field distributions of highly correlated place cells in the three different outcome groups (Sham, Recovery, No-Recovery) and the different time points (healthy, early and late after stroke). Histograms show how many place fields were located “far” from, “close” to, or “in” the reward zones of the virtual reality corridor. **G.** Analysis of the histograms in **F.** reveal that initially, highly correlated place cells in Sham mice are in particular found in reward zones, while in the late phase after stroke, animals of the Recovery group show a significant concentration of place fields close to reward zones compared to other corridor regions and compared to sham. Group differences in B-E and G were evaluated with two-way repeated measures ANOVA with the Greenhouse-Geisser correction and Tukey-Kramer multiple comparisons test. Asterisks indicate significances: *p<0.05, **p<0.01, ***p<0.001.

We then examined the location of the place fields of the synchronous active place cells for the three different experimental groups before and after stroke (Figure 5F): Although no difference of place field location was found in the healthy situation among the three groups (Figure 5G), sham animals had significantly more place fields in the reward zones than at less salient locations (far: 24.6±3.1%, in: 44.5±6.1%, p=0.029; Figure 5G). Late after stroke, animals of the Recovery group revealed a significant increase of place cell pairs with place fields close to reward zones (close: 58.8±8.8%; in: 25.2±6.9%, vs. close: p=0.003; far: 16.1±2.9%, vs. close: p<0.001; Figure 5G), indicative for a re-organization of place fields close to salient locations in animals with a good outcome, which was not detectable in mice with a chronic spatial memory deficit or Sham mice (Figure 5G).

## Discussion

We demonstrated tracking of individual neurons in the same region of interest in the hippocampus for several weeks before and after induction of microstrokes in the brain in relation to spatial memory. We used chronic 2-photon calcium imaging in mice navigating in a virtual reality corridor to dissect the functional roles of neurons in CA1 for distinct spatial information in the healthy condition and after microstrokes, and identified place cells with stable activity for a distinct place field over several days versus unstable, remapping place cells and non-coding cells for space. Furthermore, our approach allowed us to measure the loss of spatial memory: The level of cognitive decline and destruction of the neuronal network architecture in the hippocampus were strongly correlated to the number of brain-wide microstrokes identified histologically post-mortem, suggesting a dose-dependent effect, as mice with the highest load of microspheres showed the largest chronic cognitive deficit. As we found only 14% of lesions affecting directly the hippocampus and most microstrokes were remote from the recording site in the hippocampus, we not only observed a strong deficit in our spatial navigation task, but in particular an impaired stability of spatial coding of individual neurons as well as affected stability of the population activity and the persistence of the functional network structure. These results suggest that local damage to the hippocampus is not necessary to impact hippocampal function. Instead, systemic reactions to widespread microstrokes such as inflammation or network remodeling^21,22^ in form of altered input patterns might be sufficient to modify global brain activity, impairing neuronal networks remote to the acute lesion locations.

While chronic recordings of the same field of view in the hippocampus have been already reported in a few studies in the healthy condition^12,23^ and in mice after induction of epilepsy^16^ and hippocampal lesions^23^, our study shows that chronically monitoring the activity of individual cells and the same neuronal populations in the healthy condition and during several weeks after brain injury is possible. Although functional coding of individual neurons in CA1 of the hippocampus seems relatively labile in contrast to synchronously firing groups of neurons^23^ or neurons in other regions of the hippocampus^12^, we identified stable place cells which maintained their place field activity not only during the healthy condition, but also several weeks after brain-wide microstrokes. In addition, when monitoring the same neurons in sham animals for >5 weeks we found that neurons maintained their functional class (stable or unstable place cell or non-coding cell) long-term (Figure 2C, D and Figure S3), indicative for a consolidation of the functional network in sham animals (Figure 4). This is in line with preliminary results from Vaidya et al.^24^ claiming that there are two place cell pools – a transient and a sustained one – and that initially formed unstable, transient place cells are replaced by stable, sustained place cells over time. This consolidation might be mediated by behavioral timescale synaptic plasticity (BTSP), a recently discovered non-Hebbian mechanism where synaptic weights can be modulated by a single event which can be temporarily separated from the synaptic input by several seconds^25–28^ and which is also thought to induce place field formation in CA1^27^.

We found that in healthy animals with expert knowledge in the spatial navigation task place cells had a significantly higher probability to stay place cells than turning into non-coding cells. This phenomenon of “functional imprinting” was destroyed after stroke (Figure 2C, D): Instead of being pre-tuned to a distinct functional class for the spatial memory task, neurons were randomly assigned to different functions within the rewiring network after stroke. A reduction in functional imprinting after ischemic injury, as observed in this study, bears similarity to previous findings in animal models of schizophrenia^29^ and epilepsy^16^. Thus, this return to a more disordered, plastic state might not be unique to acute lesions, but also occur in chronic disruptions of hippocampal networks. It therefore may not be a purely detrimental process. Instead, it may be a cellular mechanism of increased homeostatic plasticity, which is commonly detected in the subacute phase of stroke. Besides the already known plasticity processes after brain injury such as macroanatomical map shifts^30–33^, synaptic and dendritic turn-over^34^, and molecular modifications^35,36^, the loss of functional imprinting and the promotion of neurons to new functions after stroke could be a novel cellular mechanism to better adapt to the injury and allow functional rewiring of neuronal networks after brain injury such as strokes.

While on an individual cellular level, the loss of stable coding place cells influenced the outcome (Figure 3B), on a population level microstrokes impaired location encoding (Figure 3D-F), spatial discrimination and precision (Figure 4), persistence of functional network structure (Figure 4) and synchronicity of surviving place cells (Figure 5C). All these parameters remained significantly affected in animals with a chronic deficit, while in animals with a recovery from the initial memory loss, these parameters restituted to the same level as in sham animals, highlighting these parameters as sensitive functional markers for outcome prediction and as important indicators for future interventional studies examining the positive or detrimental effects for pharmacological or rehabilitative approaches.

Finally, we found that in animals with a chronic cognitive deficit the number of highly synchronous active cells was significantly reduced. An in-depth analysis revealed that animals in the No-Recovery group lose in particular synchronously active place cells (Fig. 5C) and display nearly complete reorganization of functional connectivity networks (Figure 4G), which was not the case for the other groups. In addition, in Recovery animals place fields of synchronous active surviving place cells were in particular located close to salient locations. Thus, our results indicate a major protective mechanism: The survival of place cells which are able to maintain their place field preference for important information (e.g. salient locations such as reward zones) over time is predictive for a good cognitive outcome after stroke. Importantly, the synchronous activity of place cells helps them to survive and to re-stabilize the rewiring network after stroke. Thus, we show here that the well-known concept of Hebbian learning of “neurons that fire together wire together” applies also in a rewiring network after brain injury^36^. Elucidating this concept in the light of neuronal repair on a cellular resolution level after stroke has also implications for the development of novel pharmacological and stimulation strategies, which can induce the co-activation of neurons in networks to stabilize and enhance reorganizing circuits in the brain after stroke.

## Acknowledgements

We thank Hansjörg Kasper for technical advice and fruitful discussions as well as Antonia Weingart for support with illustrations. We thank Anna Schmidt-Rohr for programming the first GUI for tracking neurons in different experimental sessions. We thank Philipp Bethge for providing transgenic mice used in this study. This study was supported by the Branco Weiss Fellowship, the Novartis foundation for biomedical research, Dementia Research/Synapsis Foundation Switzerland, the Hurka Foundation and the TRR 274 – Checkpoints of Central Nervous System Recovery by the DFG awarded to A.S.W as well as a PhD fellowship of the Center for Neuroscience awarded to H.H.

## Autor contributions

A.S.W. ideated and provided the concept. H.H. and A.S.W. designed the study. H.H., A.S.W. and V.I. performed surgeries and carried out experiments. A.S.W., A.R. and M.W. designed and developed the virtual reality corridor. H.H., F.K., V.I., J.G. and A.S.W. performed the data analysis with input from F.H. F.H. provided resources. H.H. and A.S.W prepared figures and wrote the manuscript with input from all authors.

## Methods

### Animals

We used adult mice aged 5-7 months at the first day of experiments of both sexes (n=5 males, n=20 females). N=9 mice were C57BL/6 wild-type (Charles River, Germany) animals, n=15 were GP5.17 transgenic mice (Jackson Laboratory, RRID: IMSR\_ JAX:025393; Dana et al., 2014), and one was a triple-transgenic mouse acquired by crossing animals from the lines Snap25-IRES2-Cre-D (Jackson Laboratory, RRID: IMSR\_JAX:023525), CaMKII-tTA (Jackson Laboratory, RRID: IMSR\_JAX:007004) and Ai93D (Jackson Laboratory, RRID: IMSR\_JAX:024103). Both transgenic mouse lines expressed the calcium indicator GCaMP6f (*36*) in CA1 pyramidal neurons of the hippocampus. Mice displayed strong and uniform GCaMP6f expression in the CA1, with GP5.17 mice more readily available due to faster breeding. Mice were housed in groups of two to four under a constant 12-hour dark/light cycle, constant room temperature (22±1 °C) and with food and water ad libitum in standard cages (530 cm² floor area, 7.4 L). After starting training in the virtual reality corridor, mice were transferred to larger cages (1800 cm² floor area, 51 L) equipped with a running wheel to enhance fitness and treadmill motivation. All experiments were carried out during the active (dark) cycle of the animals and according to the guidelines of the Federal Veterinary Office of Switzerland and the license ZH241/2018 approved by the Cantonal Veterinary Office in Zurich, Switzerland. They are in accordance with the Stroke Therapy Academic Industry Roundtable (STAIR) criteria for preclinical stroke investigations (37). The samples size for the different experimental groups was estimated by means and variance of measured data in related work (*29,30,38*) and predicted to be sufficient to detect a statistically significant result in ANOVA with p<0.05 and power >0.8.

### Surgeries

For all surgical procedures except microsphere injections, mice were deeply anesthetized with 4% Isoflurane (700-800 mL O_2_ flow rate). 20-30 min prior to surgery Carprofen (5 mg/kg body weight subcutaneous (s.c.)) was administered, vitamin A crème (Bausch & Lomb) was applied to both eyes, and body temperature was maintained at 36.5 °C via a heating pad. Animals were fixed in a stereotaxic frame (Kopf Instruments) under 2% Isoflurane for surgical procedures. After surgery, mice were kept on a heating pad until being fully awake again and moving in the cage. Post-surgical pain was managed by Carprofen injections (s.c.) every 12h for 1 day, and every 24h for an additional 2 days.

### Viral injections

To induce expression of the calcium indicator GCaMP6f in wild-type mice, 300 nL of AAV9-hSyn::GCaMP6f (1×10^13^ vg/mL; Addgene, catalog # 100837-AAV9) were stereotaxically injected into the hippocampus (left CA1 at −2 mm AP, −1.3 mm ML) at least 4 weeks before the first session of two-photon calcium imaging. Injections were performed using a glass micropipette (intraMark, Blaubrand, 10-20 µm tip diameter) connected to a syringe and a stereotaxic micromanipulator (Kopf Instruments). After removing the scalp and drilling a hole above the injection site, the pipette was gradually inserted to the intended depth (−1.5 mm from skull surface), and the virus was injected with a flow rate of 0.1 µL/min. The pipette remained in place for 10 min post-injection to facilitate viral diffusion and reduce backflow before slow withdrawal.

### Chronic hippocampal imaging preparation

For performing chronic two-photon calcium imaging, we implanted a chronic cannula implant above the left CA1 region of the hippocampus 2-3 weeks after viral injection of GCaMP6f, according to published protocols (*11,39,40*). After scalp removal, iBond (iBond Total Etch, Kulzer) was applied to the cleaned skull, followed by a 3 mm craniotomy and inserting a biopsy punch above CA1 (−2 mm AP, −1.5 mm ML relative to bregma, 1.3 mm depth from skull surface). The punch was withdrawn after 10 min, and the severed tissue was slowly aspirated using a blunt 22G needle while irrigating with saline, until the corpus callosum was fully exposed. After bleeding was stopped with hemostatic sponge, a custom stainless steel cannula (3 mm diameter, 1.3 mm length) sealed with a coverslip (3 mm diameter, 0.17 mm thickness) via UV-cured dental cement (Tetric EvoFlow A1, Ivoclar Vivadent) was inserted into the brain to cover the corpus callosum and fixed to the skull with dental cement. In addition to the pain medication (Carprofen), mice received the antibiotic Baytril (10 mg/kg s.c., Bayer) for 2 days. 1 week after cannula insertion, a custom aluminium headpost (10 mm inner diameter, 12.3 mm outer diameter, 1 mm thickness) was centered on the hippocampal window and attached to the cement with a ceramic composite (Charisma, Kulzer) and additional dental cement. After UV curation, the inside wall of the cement was coated with black nail polish to minimize imaging noise. After a recovery period of 3 days, handling and behavioral training started.

### Intraarterial microsphere injection

To induce hemisphere-wide microlesions, we injected red-fluorescent 20-µm diameter PMMA microbeads (Poly-An, Cat. No. 19096-2) into the left common carotid artery (CCA) of the animals (*41,42*). Mice were deeply anesthetized with Medetomidine (250 µg/kg, Domitor, Orion Pharma), Midazolam (5 mg/kg, Dormicum, Roche) and Fentanyl (50 µg/kg, Sintetica), and positioned on their back on a heating pad. A midline incision above the thyroid gland exposed the CCA, into which 10-20 µL of the 1% microsphere stock solution, diluted in 180 µL saline was injected with a 33G needle. The external carotid artery was transiently ligated during the injection with surgical thread, and CCA flow was slightly restricted during the injection (30 s) to minimize bleeding. After removing the needle, pressure on the injection site was applied with fine cotton swabs until bleeding ceased (6-20min). The CCA was sealed with tissue adhesive (Vetbond, 3M), the incision was sutured, and the animal was placed on a heating pad until fully awake. Sham-operated mice underwent identical surgery, without the microsphere injection. After a 2-day recovery, experiments resumed on the third day.

### Virtual environment and task

We built a custom-made virtual reality (VR) setup, optimized to fit under a standard commercial two-photon microscope (HyperScope, Scientifica Ltd.). Animals could navigate in the virtual reality by walking on a custom treadmill consisting of a 5 cm wide black velvet ribbon belt stretched over two 10.5-cm diameter plastic wheels. The belt rested on a Teflon bar, with a smooth underside to reduce friction and a felted surface for improved grip. An optical rotary encoder (1440 pulses/rotation, Phidgets, Cat. No. 3530\_1) captured back wheel movements, controlling the VR displayed on three TFT-LCD monitors (10.1”, 1366x738, LG, Cat. No. LP101WH1) positioned around the animal in 11-17 cm distance to its eyes. The virtual environment created with Unity (v2018) consisted of a linear corridor featuring four distinctly patterned sections divided by reward zones (RZs) marked by salient visual and auditory (200 ms, 8 kHz, 60 dB) cues. RZs were 40 cm wide and centered at 30, 137, 243 and 350 cm in the standard 4 m corridor. We chose a multisensory environment similar to published designs (11) to improve learning rate and spatial coding in hippocampal CA1. Mice had to report RZs to receive water rewards (10 µL, controlled by a solenoid valve (SMC, Cat. No. VDW22JA)) by touching the metal spout with their tongue, where a capacitance sensor detected these touches as licks. When the animal reached the end of the corridor, it was virtually reset to the start of the corridor during a 1 s screen blackout. The VR was controlled by a Python 3.7 script, while a LabView2014 program orchestrated data acquisition and flow between the sensors, microscope and VR.

### Training protocol

At least three days after head-post implantation, the drinking water of the mice was replaced with 2% citric acid water. This method was shown to induce thirst and motivation for water rewards without restricting home-cage water access (*43*), while the body weight was controlled to be kept at >85% of the pre-experimental value. After familiarizing the mice with handling and head fixation over 2–4 days, they underwent training to selectively lick in designated reward zones (RZs) through 15 - 30-minute daily sessions. Initially, passive water rewards were given regardless of the RZ location after running 20–30 cm. After two to three sessions, passive rewards were restricted to RZ locations in a 170 cm corridor. Once mice were running consistently, the corridor length was extended to 400 cm by adjusting the gain of the rotary encoder, as well as introducing active rewards requiring licking within an RZ to receive water. Animals were not punished for licking behavior outside RZs.

Task performance was quantified per session using the spatial information (SI) content of the lick histogram. First, a lick histogram was created by binning licks (defined as the time points when the tongue of the animal touched the water spout) into 120 spatial bins, and computing the lick probability (the fraction of trials with at least one lick) for each position bin (**Figure 1**E). The SI content of this lick histogram was then determined using a formula commonly applied to measure neural spatial information content(*17,44*):

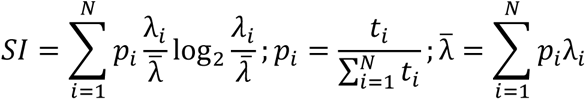

where *t_i_* represents the occupancy time spent in the *i*-th bin and λ_i_ represents the lick probability in the *i*-th bin. Animals were considered well-trained once they consistently performed at a plateau level for three consecutive days, and imaging sessions began. Animals required 13-22 days to learn the task, after which baseline neuronal activity in the CA1 was recorded across five days using two-photon calcium imaging.

### Experimental stages & groups

To capture temporal developments of behavioral and neural metrics in relation to the microsphere injections, we structured the experiment into three phases: “healthy” (before injection), “early post-stroke” (≤7 days after injection), and “late post-stroke” (>7 days – 28 days after injection). Mice that received microsphere injections and performed below 75% of their prestroke VR performance in both early and late post-stroke phases were labeled “No-Recovery”, whereas animals with an average relative VR performance of ≤75% only in the early, not late post-stroke phase, were categorized as “Recovery”. All mice with an initial performance deficit of ≤75% after microsphere injections were grouped as “Stroke”. Mice that maintained >75% relative VR performance in both post-stroke phases after microsphere injections and did not have significantly more detected spheres than sham-operated mice in their brains were pooled with sham-operated mice for subsequent analysis (**Figure S1**C). N=5 independent experiments were performed with stroke and sham animals.

### Two-photon imaging

Chronic two-photon calcium imaging of mice navigating in the VR corridor was performed with a standard commercial two-photon Galvo-Resonant scanning imaging system (HyperScope, Scientifica Ltd.) controlled by ScanImage v2017b (Vidrio Technologies; Pologruto et al., 2003). A 920-nm laser beam from a tunable Ti:Sapphire laser (Mai Tai BB, Spectra-Physics) was targeted onto CA1 pyramidal neurons through a 16× water-immersion objective (0.8 NA, Nikon). Emitted fluorescence was detected by a GaAsP photomultiplier tube (Scientifica Ltd.) after passing through a 525/50 nm band-pass filter. Beam power under the objective was adapted to GCaMP6f expression levels to 25-60 mW. The emission light path between the focal plane and the objective was shielded with a 3D-printed plastic cylinder fixed between the objective and the head-post, as well as black nail polish on the dental cement of the window preparation, to reduce light contamination from the VR monitors. Images of 512×512 pixels, corresponding to a field of view (FOV) of 830×830 µm, were acquired at a frame rate of 30 Hz. Prior to the experiment, a network with strong GCaMP6f expression encompassing a maximal number of pyramidal cells, was chosen for each mouse, and the FOV was manually aligned with this network on consecutive imaging sessions to monitor as many neurons as possible over time. Pre-processing of all two-photon imaging data was performed with CaImAn (*46*), utilizing constrained non-negative matrix factorization to perform piecewise-rigid motion correction, functional region of interest (ROI) extraction, background correction and quality evaluation. CaImAn parameters were adapted for each mouse to detect the maximum number of active cells while correctly rejecting non-neuronal ROIs, and were kept constant across the experiment. The resulting fluorescence traces were detrended with CaImAn using the following formula:

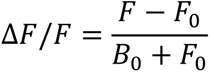

where *F* is the background-corrected fluorescence trace of the neuronal ROI, and *B*_0_ and *F*_0_ are the baseline fluorescence traces of the background and ROI respectively. The baseline of each trace is set at the percentile of the mode of the data, which was computed for each trace over a sliding window of 1000 frames using a diffusion kernel density estimator (*46,47*), which yielded baseline percentiles of 45.4±5.5 (mean ± standard deviation) across all ROIs. Finally, the resulting ΔF/F traces were transformed for spike rate analysis into deconvolved spike probabilities with the CASCADE algorithm (*48*).

### Single cell tracking

Aligning the FOV to the same network in subsequent imaging sessions enabled us to semi-automatically match individual neuronal ROIs across the experimental timeline. To identify the same cells across sessions, we computed the shifts between both FOVs using non-rigid translation, accounting for non-rigid changes in the underlying tissue, particularly post-injection, occurring over weeks. Each FOV was split into four patches, estimating the subpixel translation shift for each patch with scikit-image’s (v0.19.2) phase cross-correlation using fast Fourier transform(*49,50*), and upscaling single-patch shifts via spline interpolation to generate pixel-wise shifts. Putative matched cells were first selected by identifying the nearest neighbor for each neuron through a k-d tree (Scipy v1.10.0(*51*)), then visually inspected and curated in a custom-built interactive web application developed with Dash (v2.0.0, Plotly Technologies). Only ROIs that were manually confirmed to be the same neuron were used for single-cell analysis.

### Neuronal activity analysis

#### Linear corridor analysis

Place cell classification was performed according to published criteria (11,39). First, ΔF/F traces and occupancy times were spatially binned using 5 cm wide bins and a running velocity threshold of 5 cm/s to yield spatial activity maps for each neuron. After smoothing with a Gaussian kernel (σ = 5 cm), spatial activity maps were screened for putative place fields which were defined as locations with a ΔF/F value above 25% of the difference between the maximum and baseline ΔF/F (average ΔF/F of the lower quartile activity) of this trace. Putative place fields also had to pass three criteria: (1) the place field had to have a minimum width of 15 cm, (2) the mean ΔF/F inside the field had to be 6x higher than outside the field, and (3) significant transients had to be present for at least 20% of the time the animal moved inside the field. Significant transients were periods of at least 0.5 seconds in the unbinned ΔF/F trace with fluorescence above 3 σ (noise level σ estimated from FWHM of the ΔF/F distribution). The significance of the place field p_pf_ was estimated using bootstrapping (39). The ΔF/F trace was split into 50 frames long pieces and randomly shuffled, and place cell detection was performed on the shuffled trace. This process was repeated 1000 times, and p_pf_ was defined as the fraction of shuffles where a place field passed all three criteria. Cells were classified as a place cell if their spatial activity maps contained at least one place field that passed all three criteria and had a significance of p_pf_ ≤ 0.05.

The within-session stability of the spatial activity map of each neuron was determined by computing the Pearson correlation coefficient between the first and second halves of the trials, as well as between odd and even trials. The two coefficients were Fisher Z-transformed and averaged to derive a single stability measure (*52*).

To quantify cross-session stability of place cells, we computed the Pearson correlation coefficient between the spatial activity maps of all session pairs separated by 3 days across all tracked cells, and the Fisher transformed coefficients were averaged across all sessions within each period, yielding a stability score per phase for each neuron. The baseline stability score for a network was established as the median prestroke stability score across all neurons within that network. Subsequently, each place cell was categorized as “unstable” or “stable” for the early and late post-stroke phases based on whether its stability score for that phase was lower or higher, respectively, than baseline stability score of its network.

#### Population vector correlation

The population vector correlation (PVC) for a population of N neurons between two corridor positions *x* and *y* was defined as (*52*):

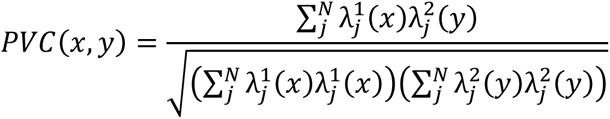

where 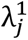 and 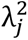 are the spatial activity maps of neuron *j* in two different sessions. Full PVC curves were created by averaging the PVC values across all corridor positions for position pairs with a Δ*x* = |*x* − *y*| location offset ranging from 0 to 275 cm. Two metrics of the PVC curves were used for further quantification: (1) The y-intercept of the curve (Δ*x* = 0 cm) represents the correlation of population activity at the same corridor position between two days, indicating the cross-session stability of the network. (2) The maximum absolute initial slope of the curve (0 ≤ Δ*x* ≤ 100 cm) represents the reduction of correlation at increasing Δ*x*, indicating the level of spatial precision of the network.

#### Bayesian decoder

A Bayesian decoder was developed to assess the predictability of corridor position based on neural activity (*52*). All models used the top 100 neurons with the highest within-session stability that were present in the training and decoding datasets. To construct the probability function for the decoder, spatial activity maps of training trials for all neurons were smoothed with a Gaussian kernel (σ = 5 cm) and the mean and standard deviation (SD) across all training trials were computed. Spatial bins with a cross-trial SD below the average per-bin SD of that neuron had their SD set to the mean SD. The estimated position of the animal 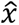 at time *t* using the neural activity of *N* neurons was defined as:

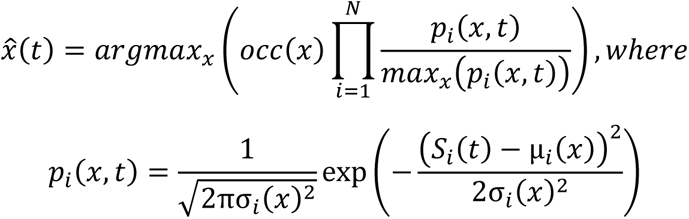

where *occ*(*x*) is the occupancy probability per spatial bin *x* in the training dataset, *μ_i_* (*x*) is the mean and *σ_i_* (*x*) is the SD of the spatial activity map at bin *x* of neuron *i* in the training dataset, and *S_i_* (*t*) is the average ΔF/F value of neuron *i* in the decoding dataset over time *t* ± Δ*t*, with Δ*t* = 0.5 s.

For within-session decoding, we adopted a leave-one-out cross-validation approach, where each trial was decoded once while the remaining trials within that session were utilized to create the training set. The final decoder performance for the session was the average metric across all iterations. For cross-session decoding, the training set was generated from the entire last pre-stroke session, and the decoding executed on the entire second session. Two different error metrics were used to quantify the decoder performance: (1) Accuracy is the fraction of correct position bin predictions out of all time points *t*. (2) Sensitivity measures the fraction of time points *t* out of all time points where the animal was inside a RZ and the decoder correctly predicted to be in a RZ. Chance levels of decoder error metrics were empirically determined by randomly shuffling the position bins in the training dataset 500 times, using the shuffled training set to decode the position of the respective trial, and averaging the error metrics of each session across all animals.

#### Functional connectivity analysis

Activity synchronicity within neuronal populations was analyzed by computing the pairwise Pearson’s correlation coefficients of the fluorescence traces of all neurons within each network. Correlation of the ΔF/F traces indicates the synchronicity of neuronal activity in time and is termed “functional connectivity”. Correlation of the spatial activity maps yields the synchronicity of neuronal activity in the linear corridor, and is termed “spatial synchronicity”. A cell pair was considered “highly synchronous” if it had a functional connectivity or spatial synchronicity value within the 95th percentile of the network.

Cross-phase changes of functional connectivity (*53*) were investigated by computing the pairwise Pearson’s correlation coefficient of ΔF/F traces for each session and averaging the coefficients of each cell pair across the sessions within each experimental phase. Neurons that were not present at all sessions during the compared pair of periods were excluded from this analysis. These phase-averaged pairwise correlation coefficients were matched across phase pairs, yielding three distributions (Healthy – Early post, Healthy – Late post, Early post – Late Post) of matched cell pairs for each mouse. For quantification, the Pearson’s correlation coefficient, corresponding p-value, and slope of a linear regression model were computed for each distributions using SciPy’s linregress function (*54*). Distributions with a p-value exceeding 0.05, indicating non-significant correlation, were excluded from the results.

Sensorimotor tasks

To assess possible deficits in the sensorimotor system after microsphere injection, animals were tested in several behavioral tasks before the injection to establish a baseline, and re-tested 2, 7, 10, 22, 32 and 35 days after microstrokes.

### Neurological deficit score

Mice were tested for neurological deficits using a common scoring system (*41,55*). In brief, possible deficits in different motor functions (limb clasping, C-shape bending, forepaw grasping and hindlimb repositioning) were scored with 0 (no deficit), 1 (moderate deficit) or 2 (severe deficit). Individual scores of each test were added to yield a total neurological deficit score.

### Skilled forelimb grasping

A grasping test was performed to assess skilled forelimb function using the MotoTrak system (*56*) (Vulintus). The setup consisted of a lever behind a acrylic glass plate, which was only accessible for the animal with one forelimb through a narrow gap in the plastic. Mice were trained to reach for the lever and pull it towards them to receive a water reward, with sessions lasting 10 – 15 min. An attempt of the animal to reach for the lever was considered a trial. If the pulling force exceeded 5 g, the trial was counted as a “hit”; if the mouse touched the lever, but without enough force, the trial was considered a “miss”; if the forelimb missed the lever, the trial was discarded. The performance per session was quantified as the hit-ratio 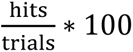.

### Open Field test

To test if microstrokes could impair mobility locomotion, mice were allowed to freely explore an empty open-roofed box of 40×40×40 cm for 6 min, once before and once in the first week after stroke. The movement of the animal was recorded with a video camera and the trajectory extracted with the EthoVision software (*57*).

### Histology

After conclusion of all experiments (6 – 8 weeks after microsphere injection), mice were deeply anesthetized with 5% Isoflurane and overdosed with pentobarbital (Kantonsapotheke Zurich, 300 mg/kg body weight, i.p. injection). As soon as respiratory arrest occurred, 0.05 mL Heparin (Braun) was injected into the left ventricle and the animal was perfused transcardially with cold 0.1M PO4 followed by 4% paraformaldehyde (PFA) in 0.1M PO_4_. Brains were extracted, post-fixed (4% PFA, 4 °C, 24 h), cryoprotected (30% sucrose, 0.1M PO_4_, 4 °C, 48 h), embedded in OCT (Tissue-Tek, Sakura), frozen at −80 °C, and 100 µm coronal sections were cut with a sliding cryostat (Microm HM 560). Free floating slices were stored in cryoprotectant (30% ethylenglycol, 15% sucrose, 0.003% Na-azide in 0.1M PO_4_). Every third slice (excluding cerebellum) was selected for immunostaining. Sections were washed 3 x 10 min in 1× PBS at room temperature (RT), incubated in blocking solution (10% natural donkey serum (NDS), 0.3% Triton X-100, 1× PBS) for 24 hours at 4 °C, followed by an incubation with guinea pig anti-GFAP (1:750, Synaptic Systems, Cat. No. 173004), rabbit anti-Iba1 (1:500, Wako, Cat. No. 019-19471) or rat anti-CD68 (FA-11 clone, 1:500, Invitrogen, Cat. No. 14-0681-82) in antibody buffer (10% NDS, 0.1% Triton X-100, 1× PBS) for 72 hours at 4 °C. Slices were washed 3 x 10 minutes in 1× PBS at RT, then incubated in secondary antibodies (Cy3-conjugated donkey anti-guinea pig, donkey anti-rabbit or donkey anti-rat, 1:250, Jackson) for 4 h at RT. Slices were exposed to DAPI (1:1000), washed again in 1× PBS and mounted on Superfrost Plus glass slides (Thermo Fisher Scientific) with fluorescent mounting medium (Dako). The whole-slice images were acquired with an Axio Scan.Z1 automatic slide scanner (10× objective, 0.6 µm/px resolution, extended depth of field; Zeiss).

### Image analysis

Images were analyzed with QuPath v0.2.3 (*58*) for microsphere and lesion detection. Experimenters were blinded to the experimental condition of each mouse, and microspheres were manually counted and assigned to brain regions using the P56 Allen Mouse Brain Reference Atlas. The non-zero sphere counts of sham-operated mice served as an estimate of the Type I error rate (false positives) of microsphere detection.

For a subset of brains, lesions around microspheres were quantified via fluorescent markers of astrocyte (GFAP) and microglia (Iba1, CD68) activation. If the average fluorescence of GFAP, Iba1, CD68 or autofluorescence (Cy5 channel, signaling cell debris) of the area surrounding a microsphere was at least 2 standard deviations higher than the same area in the contralateral hemisphere, this area was labeled as “lesioned”. Similarly, if an area not directly associated with a visible microsphere showed significant fluorescence increase compared to the contralateral hemisphere as well as control slices (slices of the same region from sham-injected animals), the area was labeled as “lesioned”. For each brain, microsphere numbers and lesion volumes were summed for each region, and the total microsphere load for the whole brain was estimated by multiplying the counts and volumes by the fraction of imaged volume of each brain based on average volumetric data provided by the Allen Mouse Brain CCFv3 (*59*).

To investigate if microsphere abundance in specific brain regions affected VR task performance, we built two generalized linear models (GLMs) using statsmodels (*60*). The dependent variables were average VR task performance during early and late post-stroke phases relative to healthy baseline performance. Independent variables were microsphere counts in a set of brain regions (hippocampus, neocortex, striatum, thalamus, white matter, other), which collectively cover the entire brain. We used a Gamma distribution for microsphere counts and the identity link function, assuming an additive effect of the number of microsphere on task performance.

### Data management and statistical analysis

Experimental data and analysis pipelines were managed by a custom DataJoint database (*61*) (RRID:SCR\_014543) implemented in Python v3.7. Statistical analysis and plotting was performed with Python and Prism v10 (Graphpad), while figures were assembled with Adobe Illustrator (v28.3). Unless stated otherwise, data are reported as mean ± standard error, and data points in figures represent individual animals, with analyses performed per session and averaged across days of each experimental phase. Boxplots are drawn with the box extending from the 25th to 75th percentiles, and the middle line plotted at the median. Whiskers reach to the minimum and maximum values of the distribution. Details regarding the statistical tests employed, multiple hypothesis correction, and the use of repeated-measures statistical testing are outlined in the figure captions and listed in Supplementary Table 1. The codes used for the processing and analysis of the raw data are available on our github page (https://github.com/Wahl-lab) on reasonable request.

### Data and materials availability

Raw and processed data are available on the data platform DANDI: https://doi.org/10.48324/dandi.001184/0.240829.1458.

All remaining data are available in the manuscript or the supplementary materials.

## References

1. Seshadri, S. & Wolf, P. A. Lifetime risk of stroke and dementia: current concepts, and estimates from the Framingham Study. Lancet Neurology Preprint at 10.1016/S1474-4422(07)70291-0 (2007).

2. Ivan, C. S. et al. Dementia after stroke: The Framingham study. Stroke (2004) doi:10.1161/01.STR.0000127810.92616.78.

3. Vermeer, S. E. et al. Silent Brain Infarcts and the Risk of Dementia and Cognitive Decline. New England Journal of Medicine 348, (2003).

4. van de Pol, L., Gertz, H. J., Scheltens, P. & Wolf, H. Hippocampal atrophy in subcortical vascular dementia. Neurodegener Dis 8, 465–469 (2011).

5. Hsu, M. & Buzsáki, G. Vulnerability of mossy fiber targets in the rat hippocampus to forebrain ischemia. Journal of Neuroscience 13, (1993).

6. Schmidt-Kastner, R. & Freund, T. F. Selective vulnerability of the hippocampus in brain ischemia. Neuroscience 40, (1991).

7. Pulsinelli, W. A., Levy, D. E. & Duffy, T. E. Regional cerebral blood flow and glucose metabolism following transient forebrain ischemia. Ann Neurol 11, (1982).

8. Johnson, A. C. Hippocampal Vascular Supply and Its Role in Vascular Cognitive Impairment. Stroke vol. 54 Preprint at 10.1161/STROKEAHA.122.038263 (2023).

9. McEwen, B. S. The plasticity of the hippocampus is the reason for its vulnerability. Seminars in the Neurosciences 6, (1994).

10. O’Keefe, J. & Dostrovsky, J. The hippocampus as a spatial map. Preliminary evidence from unit activity in the freely-moving rat. Brain Res 34, (1971).

11. Benna, M. K. & Fusi, S. Are place cells just memory cells? Memory compression leads to spatial tuning and history dependence. bioRxiv (2019) doi:10.1101/624239.

12. Hainmueller, T. & Bartos, M. Parallel emergence of stable and dynamic memory engrams in the hippocampus. Nature (2018) doi:10.1038/s41586-018-0191-2.

13. Hainmüller, T. & Bartos, M. Parallel emergence of stable and dynamic memory engrams in the hippocampus. Nature 558, 292–296 (2018).

14. Gonzalez, W. G., Zhang, H., Harutyunyan, A. & Lois, C. Persistence of neuronal representations through time and damage in the hippocampus. Science (1979) 365, 821–825 (2019).

15. Vaidya, S. P., Chitwood, R. A., Magee, J. C. & Duncan, D. The formation of an expanding memory representation in the hippocampus. bioRxiv 2023.02.01.526663 (2023) doi:10.1101/2023.02.01.526663.

16. Shuman, T. et al. Breakdown of spatial coding and interneuron synchronization in epileptic mice. Nat Neurosci 23, (2020).

17. Robinson, N. T. M. et al. Targeted Activation of Hippocampal Place Cells Drives Memory-Guided Spatial Behavior. Cell 183, 1586–1599.e10 (2020).

18. Dupret, D., O’Neill, J., Pleydell-Bouverie, B. & Csicsvari, J. The reorganization and reactivation of hippocampal maps predict spatial memory performance. Nat Neurosci 13, 995–1002 (2010).

19. Xu, H., Baracskay, P., O’Neill, J. & Csicsvari, J. Assembly Responses of Hippocampal CA1 Place Cells Predict Learned Behavior in Goal-Directed Spatial Tasks on the Radial Eight-Arm Maze. Neuron 101, 119–132.e4 (2019).

20. Wilson, M. A. & McNaughton, B. L. Dynamics of the hippocampal ensemble code for space. Science (1979) 261, (1993).

21. Anrather, J. & Iadecola, C. Inflammation and Stroke: An Overview. Neurotherapeutics vol. 13 Preprint at 10.1007/s13311-016-0483-x (2016).

22. Cirillo, C. et al. Post-stroke remodeling processes in animal models and humans. Journal of Cerebral Blood Flow and Metabolism vol. 40 Preprint at 10.1177/0271678X19882788 (2020).

23. Gonzalez, W. G., Zhang, H., Harutyunyan, A. & Lois, C. Persistence of neuronal representations through time and damage in the hippocampus. Science (1979) 365, (2019).

24. Vaidya, S. P., Chitwood, R. A., Magee, J. C. & Duncan, D. The formation of an expanding memory representation in the hippocampus. bioRxiv (2023).

25. Xiao, K., Li, Y., Chitwood, R. A. & Magee, J. C. A critical role for CaMKII in behavioral timescale synaptic plasticity in hippocampal CA1 pyramidal neurons. Sci Adv 9, (2023).

26. Priestley, J. B., Bowler, J. C., Rolotti, S. V., Fusi, S. & Losonczy, A. Signatures of rapid plasticity in hippocampal CA1 representations during novel experiences. Neuron 110, (2022).

27. Bitner, K. C. et al. Conjunctive input processing drives feature selectivity in hippocampal CA1 neurons. Nat Neurosci 18, (2015).

28. Grienberger, C. & Magee, J. C. Entorhinal cortex directs learning-related changes in CA1 representations. Nature 611, (2022).

29. Zaremba, J. D. et al. Impaired hippocampal place cell dynamics in a mouse model of the 22q11.2 deletion. Nat Neurosci 20, (2017).

30. Murphy, T. H. & Corbet, D. Plasticity during stroke recovery: From synapse to behaviour. Nature Reviews Neuroscience Preprint at 10.1038/nrn2735 (2009).

31. Wahl, A. S. et al. Optogenetically stimulating intact rat corticospinal tract post-stroke restores motor control through regionalized functional circuit formation. Nat Commun 8, (2017).

32. Wahl, A. S. et al. Asynchronous therapy restores motor control by rewiring of the rat corticospinal tract after stroke. Science (1979) 344, (2014).

33. Rehme, A. K., Fink, G. R., Von Cramon, D. Y. & Grefkes, C. The role of the contralesional motor cortex for motor recovery in the early days after stroke assessed with longitudinal fMRI. Cerebral Cortex (2011) doi:10.1093/cercor/bhq140.

34. Brown, C. E., Wong, C. & Murphy, T. H. Rapid morphologic plasticity of peri-infarct dendritic spines after focal ischemic stroke. Stroke 39, (2008).

35. Li, S. et al. An age-related sprouting transcriptome provides molecular control of axonal sprouting after stroke. Nat Neurosci (2010) doi:10.1038/nn.2674.

36. Joy, M. T. & Carmichael, S. T. Encouraging an excitable brain state: mechanisms of brain repair in stroke. Nature Reviews Neuroscience Preprint at 10.1038/s41583-020-00396-7 (2021).

37. Dana, H. et al. Thy1-GCaMP6 Transgenic Mice for Neuronal Population Imaging In Vivo. PLoS One 9, e108697 (2014).

38. Chen, T. W. et al. Ultrasensitive fluorescent proteins for imaging neuronal activity. Nature 499, 295–300 (2013).

39. Fisher, M. Recommendations for standards regarding preclinical neuroprotective and restorative drug development. Stroke 30, 2752–2758 (1999).

40. Dombeck, D. A., Harvey, C. D., Tian, L., Looger, L. L. & Tank, D. W. Functional imaging of hippocampal place cells at cellular resolution during virtual navigation. Nat Neurosci 13, 1433– 1440 (2010).

41. Pilz, G.-A. et al. Functional Imaging of Dentate Granule Cells in the Adult Mouse Hippocampus. Journal of Neuroscience 36, 7407–7414 (2016).

42. Silasi, G., She, J., Boyd, J. D., Xue, S. & Murphy, T. H. A mouse model of small-vessel disease that produces brain-wide-identified microocclusions and regionally selective neuronal injury. Journal of Cerebral Blood Flow and Metabolism 35, 734–738 (2015).

43. Balbi, M. et al. Longitudinal monitoring of mesoscopic cortical activity in a mouse model of microinfarcts reveals dissociations with behavioral and motor function. Journal of Cerebral Blood Flow and Metabolism 39, 1486–1500 (2018).

44. Urai, A. E. et al. Citric acid water as an alternative to water restriction for high-yield mouse behavior. eNeuro 8, 1–8 (2021).

45. Skaggs WE, McNaughton BL, Gothard KM, M. E. An Information-Theoretic Approach to Deciphering the Hippocampal Code. Proceedings of the IEEE (1993).

46. Pologruto, T. A., Sabatini, B. L. & Svoboda, K. ScanImage: Flexible software for operating laser scanning microscopes. Biomed Eng Online 2, 1–9 (2003).

47. Giovannucci, A. et al. CaImAn an open source tool for scalable calcium imaging data analysis. Elife 8, (2019).

48. Botev, Z. I., Grotowski, J. F. & Kroese, D. P. Kernel density estimation via diffusion. The Annals of Statistics 38, 2916–2957 (2010).

49. Rupprecht, P. et al. A database and deep learning toolbox for noise-optimized, generalized spike inference from calcium imaging. Nat Neurosci 24, 1324–1337 (2021).

50. Van Der Walt, S. et al. Scikit-image: Image processing in python. PeerJ 2014, e453 (2014).

51. Guizar-Sicairos, M., Thurman, S. T. & Fienup, J. R. Efficient subpixel image registration algorithms. Opt Lett 33, 156 (2008).

52. Virtanen, P. et al. SciPy 1.0: fundamental algorithms for scientific computing in Python. Nat Methods 17, (2020).

53. Shuman, T. et al. Breakdown of spatial coding and interneuron synchronization in epileptic mice. Nat Neurosci 23, 229–238 (2020).

54. Wu, Y. K., Hengen, K. B., Turrigiano, G. G. & Gjorgjieva, J. Homeostatic mechanisms regulate distinct aspects of cortical circuit dynamics. Proceedings of the National Academy of Sciences 117, 24514–24525 (2020).

55. Virtanen, P. et al. SciPy 1.0: fundamental algorithms for scientific computing in Python. Nat Methods 17, 261–272 (2020).

56. Bederson, J. B. et al. Rat middle cerebral artery occlusion: Evaluation of the model and development of a neurologic examination. Stroke 17, 472–476 (1986).

57. Khodaparast, N. et al. Vagus nerve stimulation during rehabilitative training improves forelimb strength following ischemic stroke. Neurobiol Dis 60, (2013).

58. Noldus, L. P. J. J., Spink, A. J. & Tegelenbosch, R. A. J. EthoVision: A versatile video tracking system for automation of behavioral experiments. Behavior Research Methods, Instruments, and Computers 33, 398–414 (2001).

59. Bankhead, P. et al. QuPath: Open source software for digital pathology image analysis. Sci Rep 7, (2017).

60. Wang, Q. et al. The Allen Mouse Brain Common Coordinate Framework: A 3D Reference Atlas. Cell 181, 936–953 (2020).

61. Seabold, S. & Perktold, J. statsmodels: Econometric and statistical modeling with python. In 9th Python in Science Conference (2010).

62. Yatsenko, D. et al. §DataJoint Elements: Data Workflows for Neurophysiology. bioRxiv (2021).

