## Supplementary Material for "Brain-wide microstrokes affect the stability of memory circuits in the hippocampus"

for

**Supplementary Table S1**

**Supplementary Figures S1-S3**

**Table S1**

| Panel | Statistical test | Post-hoc test/multiple comparisons correction |
|---|---|---|
| 1F-G | Pearson correlation & linear regression | n.a. |
| 1J-L | RM two-way ANOVA with GG | Tukey-Kramer |
| 2B | RM two-way ANOVA | Fisher’s LSD with Bonferroni |
| 2C | RM mixed-effects model with REML | Bonferroni |
| 2D | RM two-way ANOVA | Bonferroni |
| 3A | RM mixed-effects model with REML | Tukey-Kramer |
| 3B | RM two-way ANOVA with GG | Tukey-Kramer |
| 3D-F | RM two-way ANOVA with GG | Tukey-Kramer |
| 3F | Multiple one-sample t-tests | Bonferroni |
| 4B | RM two-way ANOVA with GG | Tukey-Kramer & group-wise one-way ANOVA with GG |
| 4C | RM two-way ANOVA with GG | Tukey-Kramer |
| 4G | RM two-way ANOVA with GG | Tukey-Kramer |
| 5B-E | RM two-way ANOVA with GG | Tukey-Kramer |
| 5G | RM two-way ANOVA with GG | Tukey-Kramer |
| S1C | Kruskal-Wallis | Dunn |
| S1D | Spearman correlation & linear regression | n.a. |
| S1E | RM mixed-effects model with REML | Bonferroni |
| S1F | Generalized linear model | n.a. |
| S2A | RM mixed-effects model with REML | Bonferroni |
| S2B | RM one-way ANOVA with GG | Bonferroni |
| S2C | RM two-way ANOVA | Fisher’s LSD with Bonferroni |
| S2D-E | RM two-way ANOVA with GG | Bonferroni |
| S3 | RM two-way ANOVA | Bonferroni |

**Table S2:** Statistical test and, if applicable, post-hoc test used in the figure panels. ANOVA = analysis of variance; RM = repeated measures; GG = Greenhouse-Geisser correction; REML = restricted maximum likelihood estimation.

### Figure S1

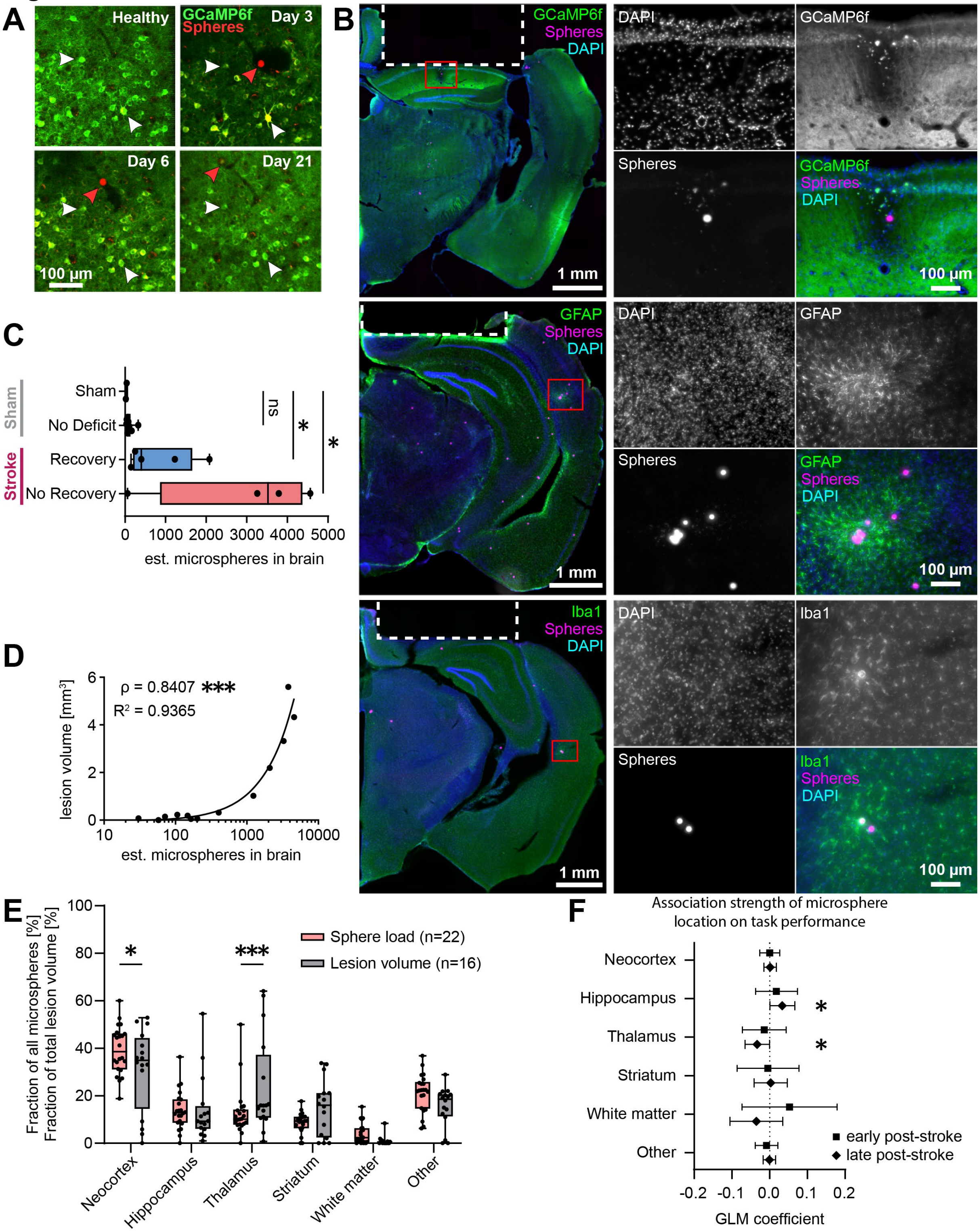

**Figure S1: Microsphere injections cause hemisphere-wide lesions. A.** Section of a field of view of a hippocampal CA1 network under the two-photon microscope at four time points. White arrows show the same neurons in each image. A red fluorescent microsphere in the tissue is visible after microsphere injection (red arrow). The black region around the sphere at day 3 is hypothesized to be microglia engulfing the foreign object and absorbing GCaMP6f fluorescence of the surrounding tissue. The sphere moves slightly with respect to the adjacent neurons over time, and does not cause visible tissue damage. **B.** Exemplary widefield images of microspheres and resulting lesions in stainings for neurons (top row, Thy-1 transgenic GCaMP6f expression), astrocytes (middle row, α-GFAP antibody), and microglia (bottom row, α-Iba1 antibody). Microspheres and lesion-caused autofluorescence is visible in the red (Cy5) channel. Left column shows whole hemispheres (scale bar = 1 mm), the right column shows high-magnification sections of microspheres and surrounding tissue (scale bar = 100 µm). Some, but not all spheres cause tissue damage, detectable by reduced GCaMP6f signal (top) as well as increased astrocyte (middle) and microglia (bottom) activity. Dashed lines in the overview images (left) indicate location of the hippocampal window implant. **C.** Extrapolated counts of microspheres in the whole brains of injected animals. Statistics assess by a Kruskal-Wallis test with Dunn's multiple comparisons test. **D.** Total lesion volume correlates strongly with number of detected microspheres. Microsphere loads show a skewed distribution with most mice having few, and few mice having many microspheres in their brains. Line shows the best fit of a linear regression model. **E.** Fraction of spheres and lesion volume across different brain regions following the Allen Atlas nomenclature. “Other” are regions not included in the other groups: cerebellum, midbrain, hindbrain, pallidum, cortical subplate, hypothalamus, olfactory areas. Statistics assessed by a subject-matched mixed-effects model with Bonferroni multiple comparisons test. **F.** Influence of microsphere counts in different brain regions on the relative VR task performance change compared to the healthy baseline performance during early and late post-stroke, evaluated by generalized linear models (Gamma distribution, identity link). Error bars show 95% confidence intervals ($n = 19$). Asterisks indicate significances: *$p<0.05$, **$p<0.01$, ***$p<0.001$.

### Figure S2

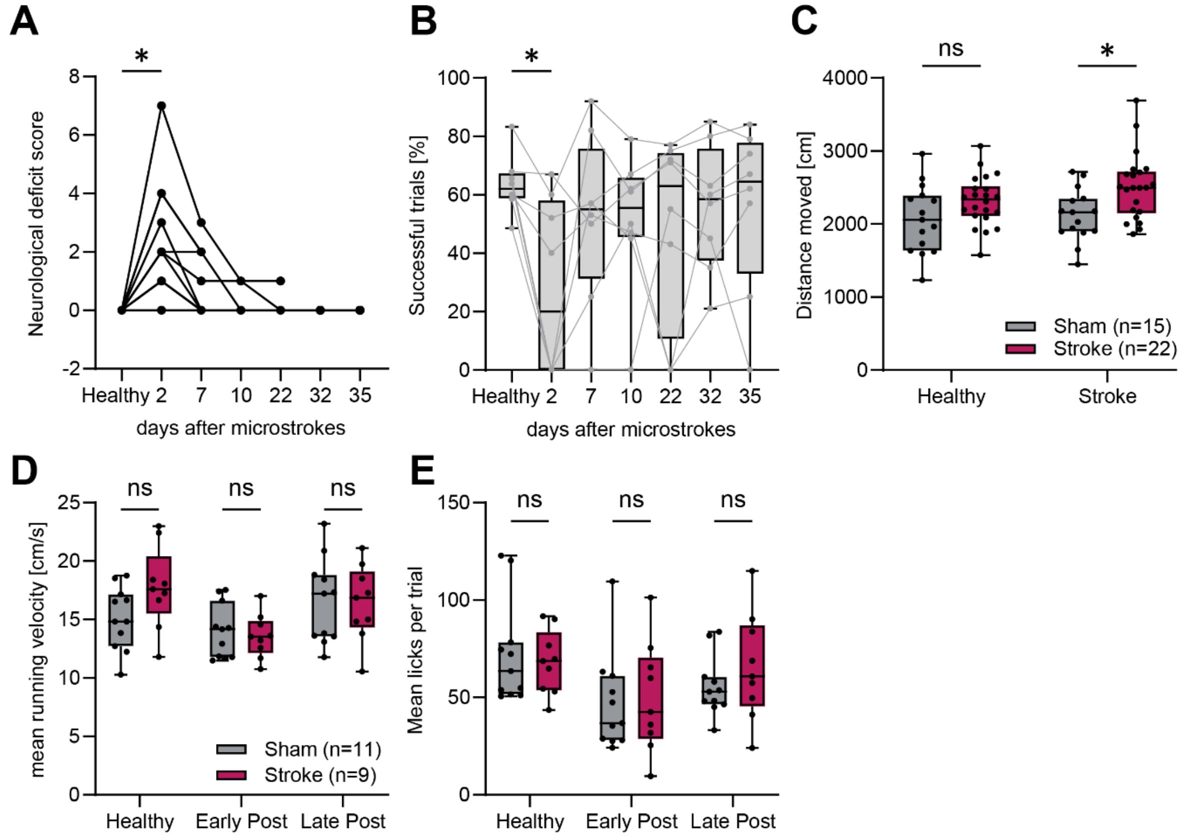

**Figure S2: Motor functions remain largely unaffected by microstrokes. A.** Neurological deficit scores[56] measured before and after microstrokes. Mice showed a significant deficit only on the first session two days after microsphere injection. **B.** Percentage of successful trials in a forelimb motor task testing strength and grasping skills measured before and after microstrokes. Mice showed a significant deficit only on the first session two days after microstrokes. **C.** Total distance moved in an open field of stroke and sham mice, tested before and within the first week after microstroke surgery. Stroke mice were more active than sham mice in the post-stroke session (Sham: 2472±92 cm, Stroke: 2506±96 cm, p=0.006). **D.** Mean running velocity of mice in the VR corridor during the experiment. Microstrokes did not affect running speed, as sham and stroke mice did not show significant differences before or after surgery. **E.** Mean number of licks per trial in the VR corridor during the experiment. Microstrokes did not affect lick behavior, as sham and stroke mice did not show significant differences before or after surgery. Statistics were evaluated using two-way repeated-measures ANOVA with Greenhouse-Geisser correction and Tukey-Kramer multiple comparisons test. Asterisks indicate significances: *p<0.05, **p<0.01, ***p<0.001.

**Figure S3**

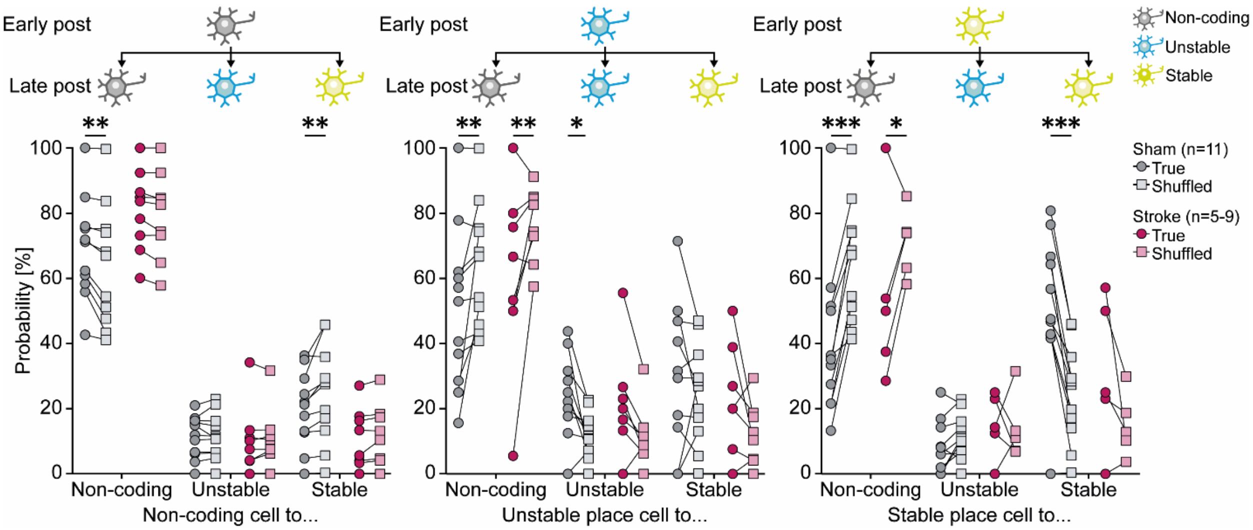

**Figure S3: Microstroke-induced loss of functional imprinting for stable place cells is partially reversed during late post-stroke.** Observed probabilities of non-coding cells (grey), unstable (blue) and stable (yellow) place cells to maintain or switch their functional class from the early to late post-stroke phase, compared against shuffled distributions. Two-way repeated-measures ANOVA with Bonferroni multiple comparisons test. Asterisks indicate significances: *p<0.05, **p<0.01, ***p<0.001.
